# Setdb1 safeguards proper differentiation of adult intestinal stem cells by controlling chromatin accessibility and transcriptome variability

**DOI:** 10.1101/2024.10.18.619079

**Authors:** Ioanna Peraki, Ioannis K. Deligiannis, Dimitris Botskaris, Marianna Stagaki, Haroula Kontaki, Elena Deligianni, Giannis Giannoulakis, Matthieu D. Lavigne, Celia P. Martinez-Jimenez, Iannis Talianidis

**Affiliations:** Institute of Molecular Biology & Biotechnology, Foundation for Research and Technology, Hellas (IMBB-FORTH), Heraklion, Crete, Greece; Dept. of Biology University of Crete, Heraklion, Crete Greece; Helmholtz Pioneer Campus (HPC), Helmholtz Zentrum Munchen, Neuherberg, Germany; TUM School of Medicine, Technical University of Munich, Munich, Germany; Institute of Biotechnology and Biomedicine (BIOTECMED), Department of Biochemistry and Molecular Biology, University of Valencia, Burjassot, Spain

## Abstract

The histone methylase Setdb1 plays a pivotal role in embryonic stem cell maintenance and developmental lineage specification. However, its function in adult stem cells remains elusive. Here we show that conditional inactivation of *Setdb1* in Lgr5^+^ intestinal stem cells alters the transcriptional programs of the progeny cell types and results in increased cell-to-cell transcriptional variability. Loss of Setdb1 blocked differentiation towards the absorptive enterocyte lineage, while the secretory cell types were only marginally affected due to the activation of alternative developmental trajectories. *Setdb1* inactivation did not affect global H3K9 methylation at large heterochromatin domains but led to altered distribution of transposase-accessible chromatin regions, aberrant exposure of transcription factor binding sites and premature activation of differentiation-specific genes. The results demonstrate that Setdb1 regulates intestinal stem cell differentiation by fine-tuning chromatin accessibility in open euchromatin regions thereby controlling transcriptional variability between cells.

## Introduction

A significant proportion of the eukaryotic genome is organized in highly compacted structures, called “constitutive” heterochromatin, which plays fundamental role in maintaining genome stability via inhibiting the activity of mobile transposable elements, silencing repetitive and retroviral sequences and ensuring proper chromosome segregation.^1–3^ Heterochromatin organization is also detectable in specific gene areas, forming locally closed but reversible structures, called “facultative” heterochromatin, which contributes to condition-specific transcriptional repression.^4^ Nucleosomes at heterochromatin are densely modified by methylations of histone 3 (H3K9Me_]_and H3K27Me_]_) or histone 4 (H4K20Me_]_)^2^. H3K9 di-and trimethylation at major blocks of repeat-rich heterochromatin, such as centromeric and telomeric regions is catalyzed by three major methyltransferases, Suv39h1, Suv39h2 and Setdb1.^5–8^

Dynamic changes in heterochromatin play crucial roles in cell lineage specification during early embryonic development, stem cell maintenance and differentiation.^9–13^ The function of the three main H3K9 methylating enzymes in these processes display considerable redundancy, as the full eradication of H3K9Me_]_marks can only be achieved in double and triple knockout cells.^5,7,9,14^ Despite this redundancy genetic inactivation of the individual enzymes often leads to aberrant phenotypes, such as impaired differentiation, genome instability, viral mimicry, necroptosis or cancer.^3,10,15^

Setdb1 has been postulated to influence gene expression via multiple mechanisms. These include control of constitutive heterochromatin spreading, formation of facultative heterochromatin,^6^ silencing LTR retrotransposon insertions^5^ and methylation of transcription factors and signaling regulators.^16–18^ Setdb1 is also important for the formation of poised enhancers, where it cooperates with NSD1 methylase to create nucleosomes with dual histone modification marks, H3K9_]_/H3K36_]_.^19^

The role of Setdb1 in embryonic stem cells (ESc) has been extensively studied.^6,12,13,20^ Apart from its pivotal role in maintaining genome stability via silencing endogenous retroviral elements,^3^ Setdb1 has been shown to cooperate with PRC2 to suppress developmental regulators^20^ and contribute to the transition of facultative heterochromatin to constitutive heterochromatin during differentiation. Consistent with this, loss of Setdb1 in embryonic stem cells and in more differentiated cells, often leads to altered transcription programs, suggesting that Setdb1-mediated silencing is important for the regulation of tissue-specific gene expression patterns.^10,12^

Multipotency of adult stem cells, similar to the maintenance of pluripotency in ES cells, is governed by the function of transcriptional regulators, which activate an array of genes characteristic to the stem cell phenotype and in parallel repress genes conferring the differentiated cell phenotype.^21^ Little is known however, how chromatin modifiers regulate multipotent phenotypes in different adult tissues.

To explore this question, we investigated the function of Setdb1 in Lgr5^+^ intestinal stem cells. Lgr5^+^ stem cells are found at the base of the intestinal crypts, embedded into a microenvironment of specialized cells, which provide the necessary repertoire of signaling molecules, like Wnt, Notch, BMP, to regulate self-renewal and differentiation.^22^ Lgr5^+^ stem cells give rise to proliferating progenitors or transit amplifying (TA) cells, that subsequently differentiate to each principal cell types of the intestinal epithelium, including Paneth cells, Goblet cells, enterocytes, enteroendocrine cells and Tuft cells.^22,23^

In this study, using lineage tracing and single-cell RNA-sequencing (scRNA-seq) combined with single-cell ATAC-sequencing (scATAC-seq) approaches, we demonstrate that Setdb1 is required for the proper differentiation of Lgr5^+^ stem cells into distinct intestinal epithelial cell types. The results reveal an unexpected function of Setdb1 in open chromatin domains, which contributes to the tight control of the expression of linked genes, which play fundamental roles in shaping cell identity in the rapidly renewing intestinal epithelium.

## Results

### *Setdb1* inactivation results in defective differentiation of Lgr5^+^ cells in the intestinal epithelium

To investigate the function of Setdb1 in the self-renewal and differentiation properties of the intestinal Lgr5^+^ adult stem cells, we crossed mice carrying Lgr5-GFP-Cre^ERT2^ knock-in allele^22^ with mice, carrying floxed exon-3 *Setdb1* alleles (Figure S1A and S1B). In these mice, Lgr5^+^ intestinal stem cells are labeled by GFP and upon tamoxifen treatment, *Setdb1* is selectively inactivated in GFP-expressing Lgr5^+^ stem cells (Figure S1E and S1F). To follow the fate of Lgr5^+^ cells during differentiation, we crossed Lgr5-GFP-Cre^ERT2^ and Lgr5-GFP-Cre^ERT2^/ Setdb1^lox/lox^ mice with ROSA (CAG-tdTomato*, EGFP*)Ees mice (abbreviated as nTnG), carrying the dual reporter of tomato-red and EGFP in the ROSA locus (Figure S1C and S1D). In these mice, after tamoxifen treatment, GFP is expressed in both, Lgr5^+^ stem cells and their progeny, due to the irreversible loss of the tdTomato gene. As expected, GFP-labeled cells were localized only in the bottom of the crypts in mock-treated cells (Figure 1A). In agreement with previous reports, which showed that Lgr5^+^ cells can give rise to all intestinal epithelial cell types,^22^ we observed GFP-labeled cells over the entire crypt and villi in a continuous fashion five days after tamoxifen treatment (Figure 1A). The labeled strip of epithelial cells originating from the bottom of the crypts was missing in some areas, which is explained by the known mosaic expression of the Lgr5 knock-in allele caused by random silencing.^22,23^

**Figure 1.**
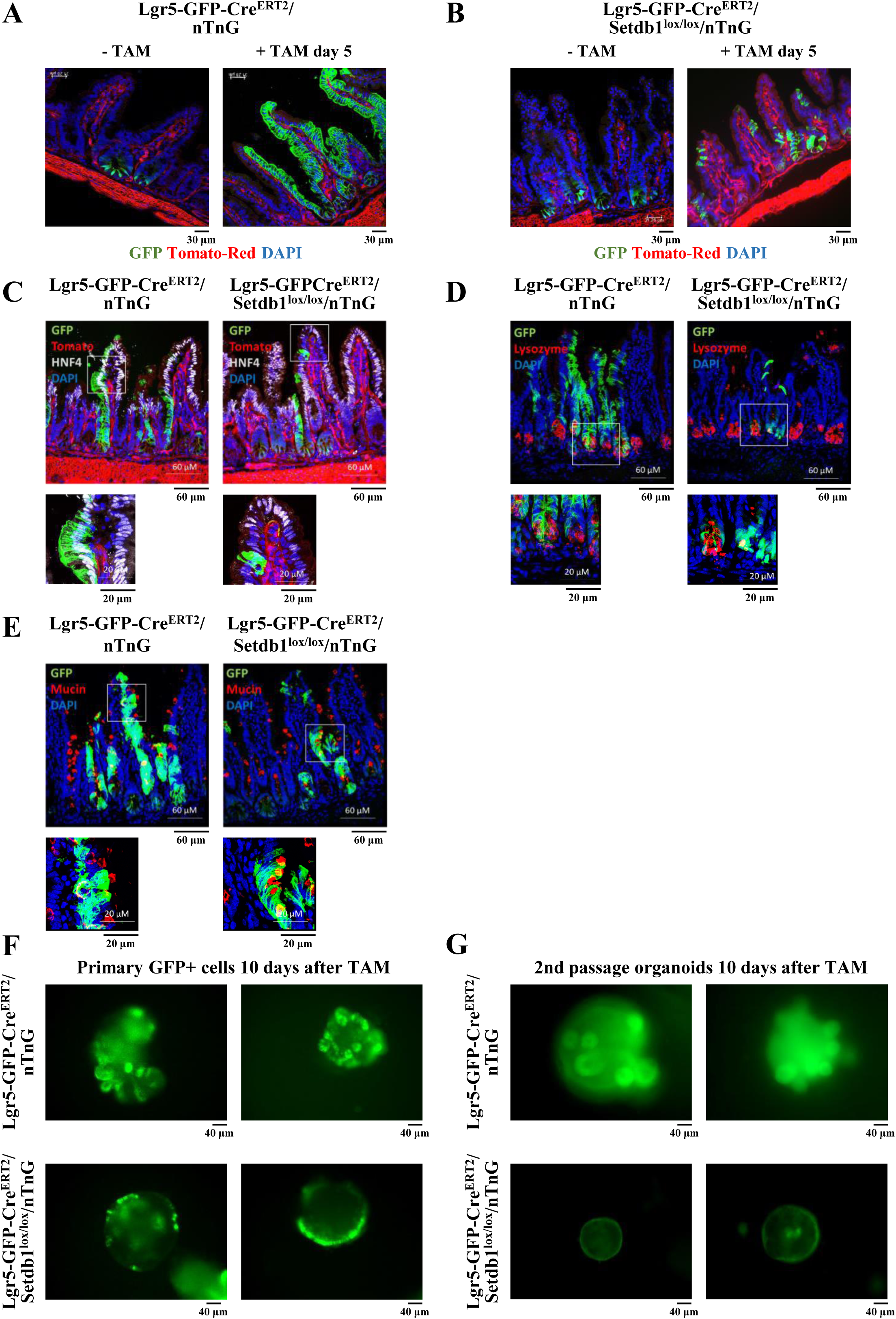
Setdb1 is required for the differentiation of Lgr5^+^ stem cells to intestinal epithelial cell types. (A-B) Representative fluorescent images from the ileum of Lgr5-GFP-Cre^ERT2^/nTnG mice, expressing wild type Setdb1 mRNA (A), and Lgr5-GFP-Cre^ERT2^/Setdb1^lox/lox^/nTnG mice, carrying floxed alleles of Setdb1 (B) that were treated with tamoxifen (+TAM) or with solvent (-TAM) for 5 days. (C-E) Representative immunofluorescence staining of ileal epithelium with antibodies recognizing the mature Enterocyte marker HNF4α (C), the Paneth cell marker Lysozyme (D) and the Goblet cell marker Mucin (E). Bottom panels correspond to zoom-in images of the indicated areas. (F) Representative images of intestinal organoids from FACS-sorted GFP^+^ cells, isolated from Lgr5-GFP-Cre^ERT2^/nTnG mice (upper panel) and Lgr5-GFP-Cre^ERT2^/Setdb1^lox/lox^/nTnG mice (bottom panel). After seeding, the cells were treated with 100nM tamoxifen (TAM) for 10 days, before image acquisition. (G) Representative images of 2^nd^ passage organoids originating from tamoxifen-treated Lgr5-GFP-Cre^ERT2^/nTnG organoids (upper panel) and from Lgr5-GFP-Cre^ERT2^/ Setdb1^lox/lox^/nTnG organoids (bottom panel). Images were taken 10 days after reseeding the cells.

In contrast, *Setdb1* inactivation in Lgr5^+^ cells resulted in only few GFP-labeled cells with scattered distribution along the crypt-villus axis and some accumulation of the labeled cells at the bottom of the crypts (Figure 1B). Double-immunostaining assays with antibodies against the enterocyte marker HNF4α, the Paneth cell marker Lyzozyme (Lyz) and the Goblet cell marker Mucin (Muc), revealed strong reduction of the HNF4α/GFP double positive cells in Setdb1-deficient mice (Figure 1C and S1G). Lyz/GFP double positive cells were reduced to a lesser degree (Figure1D and S1G) while Muc/GFP double-positive cells have almost doubled (Figure 1E and S1G). The above staining patterns suggest that Setdb1-deficiency leads to a major defect of Lgr5^+^ stem cell differentiation towards epithelial lineages, especially to enterocytes.

Independent evidence for the requirement of Setdb1 in Lgr5^+^ stem cell differentiation was provided by *in vitro* organoid formation assays. Previous studies have established that single Lgr5^+^ stem cells can differentiate *in vitro* and give rise to self-organizing organoid structures, which contain all the intestinal epithelial cell types.^24^ We isolated GFP-expressing cells from the ileum of Lgr5-GFP-Cre^ERT2^-nTnG and Lgr5-GFP-Cre^ERT2^/Setdb1^lox/lox^/nTnG mice by FACS-sorting and cultured them in organoid culture conditions. From the day of seeding, the cells were treated with tamoxifen and the formation of GFP-labeled 3D organoids was monitored. As shown in Figure 1F, 10-days after tamoxifen treatment complex 3D organoid structures were developed from Lgr5-GFP-Cre^ERT2^-nTnG cells, with the majority of the cells being GFP positive. In contrast, cells from the Lgr5-GFP-Cre^ERT2^/Setdb1^lox/lox^/nTnG mice, developed more simple structures and the cells displayed heterogeneous GFP-label intensity (Figure 1F). Importantly, when the cells from the GFP-labeled organoids were subjected to a second passage, only the cultures derived from normal *Setdb1* allele-containing stem cells gave rise to fully developed 3D GFP-labeled organoid structures. Organoids derived from Setdb1-deficient cells generated small cellular assemblies with spherical morphology and near background-level GFP fluorescence (Figure 1G).

### Setdb1 requirement for proper intestinal epithelial cell specification reveals alternative unconventional trajectories towards secretory and endocrine-related cell lineages

To determine whether Setdb1 affects Lgr5^+^ stem cell differentiation at early or late stages, we performed single-cell RNA-seq experiments after FACS sorting of GFP^+^ intestinal epithelial cells from Lgr5-GFP-Cre^ERT2^ and Lgr5-GFP-Cre^ERT2^/Setdb1^lox/lox^ mice (hereby named Lgr5^+^WT and Lgr5^+^Setdb1KO, respectively), five days after tamoxifen treatment. From two independent biological replicates, a total of 10,598 Lgr5^+^WT cells and 13,702 Lgr5^+^Setdb1KO high-quality cells with an average of 2550 genes per cell were analyzed. After data merging and integration, unsupervised graph clustering was used to partition the cells into 15 clusters and the cell types were annotated using previously described marker gene signatures.^25,26^ The results were visualized by Uniform Manifold Approximation and Projection (UMAP) dimension reduction technique (Figure S2A and S2B). About 8752 cells, corresponding to 82% of the total, expressed GFP mRNA. These cells were identified as Lgr5^+^ stem cells (Stem-I-II-III), early progenitors including TA cells (Progenitor-I-II-S) and potential lineage-specified progenitors (Enterocyte immature, Goblet-I and EEC-I). The remaining 18% of the cells corresponded to mature enterocytes, Goblet cells, Paneth cells, Tuft cells and enteroendocrine cells, whose presence is explained by the gating strategy used and the not complete enrichment of Lgr5^+^GFP^+^ cell population by our FACS sorting-mediated isolation.

We have included the latter “contaminating” cell clusters in our comparative transcriptome analyses, as they provided useful information for investigating Setdb1’s role in transcriptional transitions that define lineage trajectories during differentiation.

Regarding the stem cell compartment, each of the three distinct stem cell populations, expressed pan-stem cell markers (Figure S3A, S3B and S3C) along with their corresponding cluster-specific gene signatures. The transcriptome of Stem-I cells is characteristic to the low cycling Lgr5^+^ stem cells, while Stem-II cluster, corresponds to the recently described MHCII expressing antigen presenting stem cell population.^27,28^ Stem-III cells represent a more proliferative cell population (Figure S3A, S3B and S3C).

The transcriptome profiles of the cells termed Progenitors revealed two main distinct cell clusters. These cells correspond to the more proliferative downstream cell types including +4 cells and TA cells,^26,29^ which correspond to precursors for secretory lineages (Progenitor-I) and for enterocytes (Progenitor-II) (Figure S3D and S3E). Furthermore, transcription profile analysis resulted in the separation of three Goblet cell populations (Figure S4A, S4B and S4C) and two Enteroendocrine (EEC) cell populations (Figure S4D and S4E). The transcriptome of Goblet-I and EEC-I cells suggest that they represent lineage-specified precursors of the more mature Goblet-II and III or EEC-II cells (Figure S4). ^26,30^

To delineate the differentiation dynamics of the intestinal epithelial cell types we performed pseudotemporal reconstruction of the transcriptional changes^31,32^ and mapped seven distinct continuous trajectories in cells from Lgr5-GFP-Cre^ERT2^ (control) mice (Figure 2A and 2B). As expected, each of the seven lineages originating from Stem-I cells gave rise to all known intestinal epithelial cell types. The intermediate cellular states in the different lineages revealed interesting cell fate dynamics. Lineage-1 represents a Stem-I cell self-renewal path through Stem-II, Stem-III and Progenitor-I cellular states. Lineage-2 is the sole differentiation trajectory for the generation of mature enterocytes via Stem-II, Stem-III, Progenitor-II and Immature Enterocyte cellular states. The hierarchical path from Stem-I to Stem-II, Stem-III and Progenitor-I cells is common in Lineages 3, 4, 5, 6 and 7, which end up in Goblet cells (Lineages 3 and 4), in Paneth cells (Lineage-5) and Enteroendocrine cells (Lineages 6-7). Interestingly, for Lineage-5 that gives rise to Paneth cells, the cell trajectory analysis reconstructs a differentiation path connecting Paneth cells back to the Stem-I cluster, supporting the possibility that de-differentiation of Paneth cells may also be involved in Stem-I self-renewal, as indicated in a previous report.^33^ Similarly, the trajectory topology for Lineages 6 and 7 shows Enteroendocrine clusters as the final differentiation stage, placing the Tuft cell cluster in an intermediate stage within the differentiation process. These results indicate that Tuft cell clusters might represent an upstream and hierarchical connection between the processes regulating the differentiation of these cell types.

**Figure 2.**
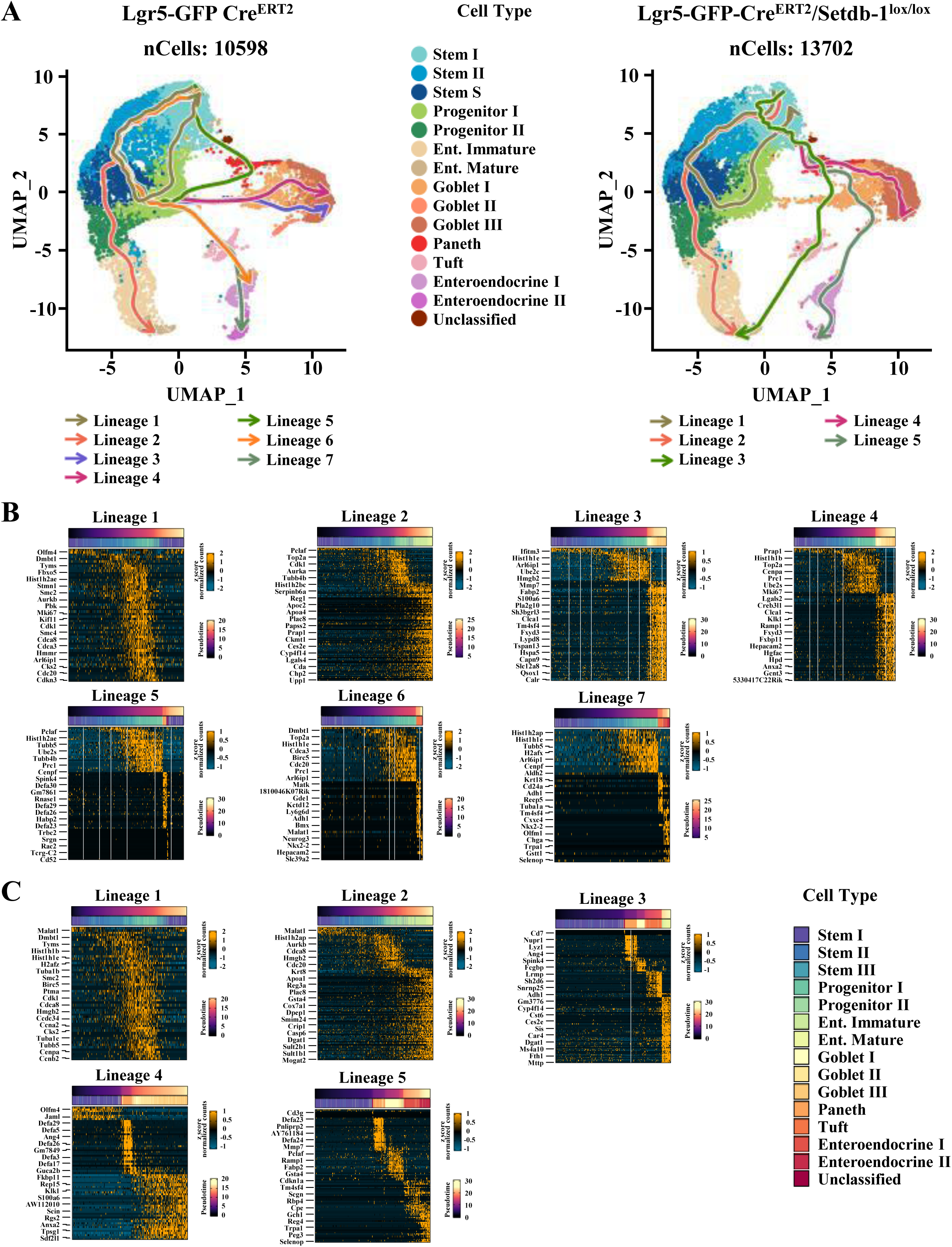
Dynamic transitions and alternative differentiation trajectories of epithelial cell types in Setdb1-deficient cells (A) UMAP projection of RNA velocities in single cells from Lgr5-GFP-Cre^ERT2^ (left panel) and Lgr5-GFP-Cre^ERT2^/ Setdb1^lox/lox^ (right panel) mice after 5 days of tamoxifen treatment. Arrows indicate lineage trajectories from pseudotemporal inference, starting from Stem-I cell cluster. The color codes corresponding to the different cell clusters are indicated at the middle. (B-C) Gene expression heatmap of differentially expressed genes of the displayed cell lineages plotted along velocity-based pseudotime starting from Stem-I cell cluster. Panels in (B) show data from Lgr5-GFP-Cre^ERT2^ mice expressing normal allele of Setdb1, while panels in (C) show data from tamoxifen treated Lgr5-GFP-Cre^ERT2^/Setdb1^lox/lox^ mice, expressing mutant allele of Setdb1. Top bars above the heatmaps indicate pseudotemporal axis. The bars below indicate corresponding cell identity across the inferred trajectory, according to the color code. Color bars at the right show z-score normalized relative mRNA read counts (upper bar) and relative pseudotimes (lower bars).

Analyses of Setdb1-deficient cells from Lgr5-GFP-Cre^ERT2^/Setdb1^lox/lox^ mice revealed major alterations of the above trajectories (Figure 2A and 2C). While Lineage-1 and Lineage-2, specifying the Stem-I self-renewal and enterocyte differentiation paths were not altered, the other trajectories specifying secretory cell types (Lineages 3, 4, 5, 6, 7) were not detected (Figure 2A and 2C). In contrast, three new lineages with aberrantly ordered paths appeared, connecting Stem-I cells directly to Paneth cells and branching either towards Tuft cells and enterocytes through Goblet-I cells (KO-Lineage 3), or towards EEC-I and EEC-II cells through Goblet-II cell cluster (KO-Lineage-5), or towards Goblet-III cells through Goblet-II cell cluster (KO-Lineage-4).

While transcriptome-based trajectories accurately recapitulate the order of the consecutive differentiation stages^31^, they do not provide quantitative and comparative assessment about the actual extent of the operation of the stable or altered differentiation paths in specific conditions. Therefore, we further analyzed the functional importance of the above trajectories by performing single-cell transcriptome analyses in lineage-traced Lgr5^+^ cells, taking advantage of the fact that following tamoxifen treatment of Lgr5-GFP-Cre^ERT2^/nTnG and Lgr5-GFP-Cre^ERT2^-nTnG/Setdb1^lox/lox^/nTnG mice, only Lgr5^+^ cell-derived descendant cell populations acquire GFP expression, which can be isolated by FACS sorting. In agreement with the immunostaining results in Figure 1, five days after tamoxifen treatment, we could detect all the known intestinal epithelial cell types in the control Lgr5-GFP-Cre^ERT2^-nTnG mice (Figure 3A). In contrast, in Setdb1-deficient cells from Lgr5-GFP-Cre^ERT2^-nTnG/Setdb1^lox/lox^ mice a dramatic accumulation of Stem-I cells was observed along with a major decrease in the number of mature enterocytes (Figure 3A, 3B and 3C). A moderate increase in the number of Goblet, Tuft and EEC cells was also observed, with the precursor Goblet-I population increasing the most. The number of Paneth cells in the Setdb1-deficient Lgr5^+^ descendants was nearly halved.

**Figure 3.**
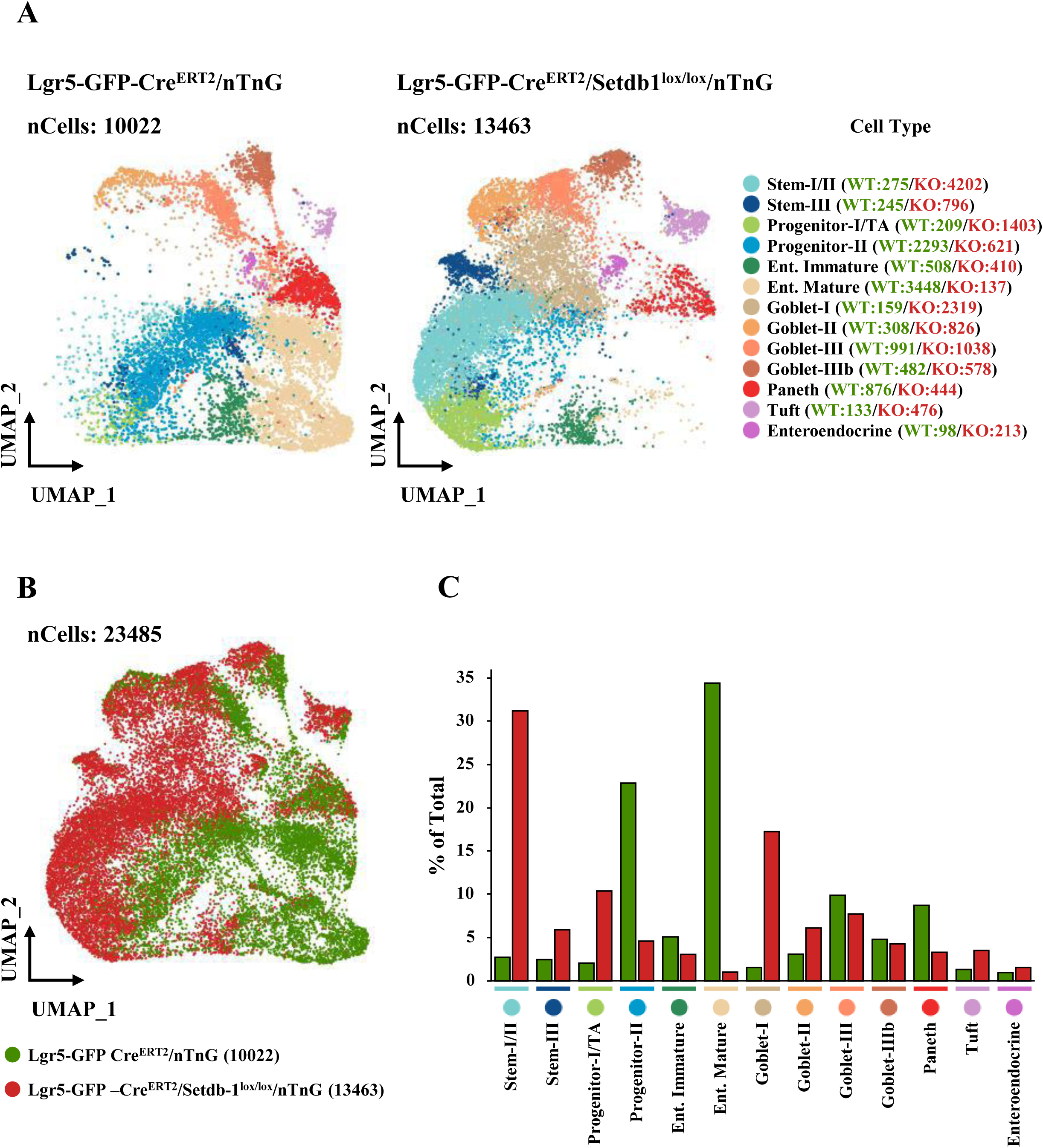
Lineage tracing of individual Lgr5^+^ progeny epithelial cells reveals defective differentiation of Setdb1-deficient Lgr5^+^ stem cells. (A) UMAP projections of cell clusters obtained from FACS-sorted GFP^+^ cells isolated from tamoxifen-treated Lgr5-GFP-Cre^ERT2^/nTnG (left panel) and Lgr5-GFP-Cre^ERT2^/ Setdb1^lox/lox^/nTnG mice (right panel). The color codes and the number of cells identified in the different cell clusters are indicated at the right. (B) Aligned UMAP plots of cells detected in samples from tamoxifen-treated Lgr5-GFP-Cre^ERT2^/nTnG (green dots) and Lgr5-GFP-Cre^ERT2^/ Setdb1^lox/lox^/nTnG (red dots) mice. (C) Comparison of the percentages of cells corresponding to the different clusters in tamoxifen-treated Lgr5-GFP-Cre^ERT2^/nTnG (green columns) and Lgr5-GFP-Cre^ERT2^/Setdb1^lox/lox^/nTnG (red columns) mice.

Taken together, the above results demonstrate that Setdb1 deficiency results in major block of the canonical pathway of Lgr5^+^ stem cell differentiation from the very early steps. Setdb1 function is fully required for enterocyte differentiation and for the progression of Progenitor-I cell population into other secretory and endocrine-related cell types. Nevertheless, these latter cell types can be generated directly from Lgr5^+^ Stem-I cells, via the activation of alternative, non-canonical differentiation trajectories.

### Setdb1 safeguards proper differentiation of Lgr5^+^ stem cells by preventing uncontrolled expansion of open accessible chromatin domains and premature aberrant gene activation

Previous studies using Villin-Cre^ERT2^ transgene, which inactivates *Setdb1* in all intestinal epithelial cell types, have demonstrated a noticeable induction of endogenous retrovirus expression, which triggered a viral mimicry and activated antiviral immune responses, culminating into inflammation and necroptosis.^34,35^ In our experimental setup, where *Setdb1* inactivation is restricted to Lgr5^+^ stem cell population and the analyses were performed as early as 5 days after Cre transgene activation, we failed to detect massive increase of retroviral transcripts by bulk RNA-seq assay or the activation of innate immunity-related or necroptosis-related genes in a substantial number of cells by scRNA-seq (Figure S5A and S5B). Although retroviral silencing is likely to occur at later time points, the above result suggests that the observed early differentiation defect is rather the result of a direct function of Setdb1 in the regulation of differentiation-specific genes at early stem cell states.

Indeed, single-cell Assay for Transposase-Accessible Chromatin sequencing (scATAC-seq) analyses identified a large number of new peaks and also a great number of peaks that were gained or lost in Setdb1-deficient Stem-I cells compared to wild-type, control cells (Fig. 4A and 4E). In addition, scATAC-seq reads near the transcription start sites (TSS) were significantly enriched in *Setdb1*-KO Stem-I cells (Figure 4B). Similarly, peak-to-gene link analysis revealed consistently more genes with ATAC-seq peak link counts in *Setdb1*-KO cells compared to wild-type control (Figure 4C and 4D), implying that more open regulatory regions are connected to target genes as a result of Setdb1 inactivation. Co-clustering analyses of the regions linked to genes in *Setdb1*-KO and wild type Stem-I cells (based on scATAC-seq and scRNA-seq signals), identified 6 major peak-to-gene clusters with significantly altered accessibility profiles and concomitant changes in gene expression (Figure 5A). Aberrantly accessible regions and linked activated genes in these clusters, included genes that are normally expressed in other differentiated intestinal epithelial cell types, such as the enterocyte-specific Rbp2, ApoC3, Papss2 genes, the Goblet cell-specific Tff3, Spink4, Mok genes, the Paneth cell specific Lyz1, Itln1 and several defensin genes, the Tuft cell specific Fes gene and a number of Entero-endocrine cell-specific genes, like Gpx3, Itg9a, Pax4, Shc2, Slc8a1, Ambp (Figure 5B). These data point to a partial loss (cluster 5) and gain (especially cluster 1 and 3) of chromatin accessibility control in Setdb1-deficient cells, which leads to aberrant and premature activation of differentiation stage specific genes in Stem-I cells.

**Figure 4.**
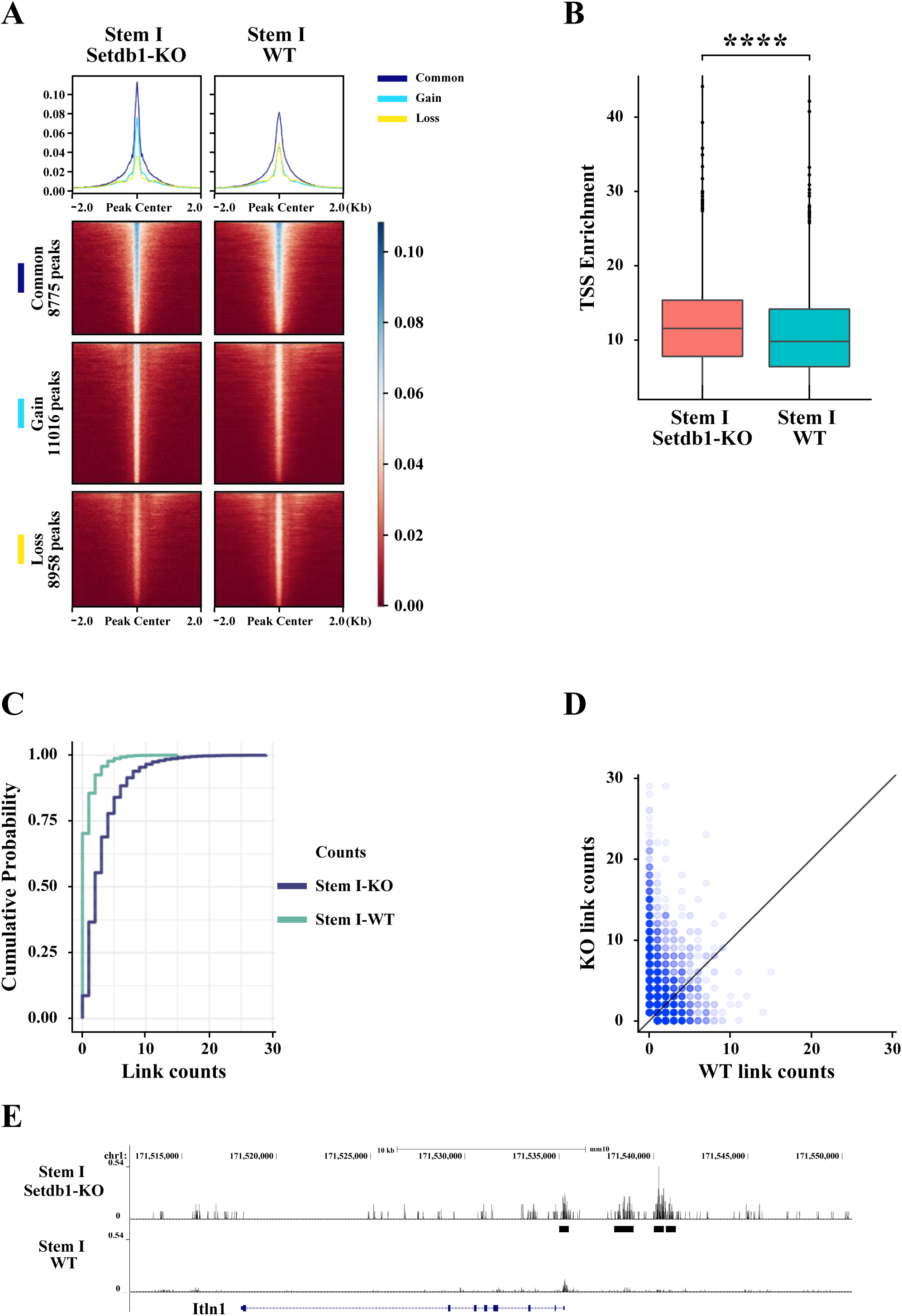
Reorganization of transposase accessible chromatin areas in Setdb1-deficient intestinal stem cells. (A) Signal intensity heatmaps and average distribution profiles (top panels) of scATAC-seq peaks in the Stem-I cell cluster from tamoxifen-treated Lgr5-GFP-Cre^ERT2^ (WT) mice and Lgr5-GFP-Cre^ERT2^/Setdb1^lox/lox^ (Setdb1-KO) mice. Read distribution at the indicated regions around the center of ATAC-seq peaks is shown. ATAC-seq peaks were clustered according to their presence in both wild type and Setdb1-KO cells (Common peaks), in Setdb1-KO cells only (Gained peaks) and in wild type cells only (Lost peaks). (B) Box plot analysis of TSS enrichment of scATAC-seq peaks in the Stem I cell cluster from tamoxifen-treated Lgr5-GFP-Cre^ERT2^ (WT) mice and Lgr5-GFP-Cre^ERT2^/Setdb1^lox/lox^ (Setdb1-KO) mice. ****p<1.116e^-11^ as determined by Wilcoxon rank sum test. (C) Quantitative comparisons of scATAC-seq peak variations in the Stem-1 cell cluster of tamoxifen-treated Lgr5-GFP-Cre^ERT2^ (WT) mice and Lgr5-GFP-Cre^ERT2^/Setdb1^lox/lox^ (Setdb1-KO) mice. The graph shows the number of ATAC-seq peaks linked-to-genes and the cumulative probability of their occurrence. (D) Scatter plot presentation of the peak-to-gene link counts in the Stem-1 cell cluster of tamoxifen-treated Lgr5-GFP-Cre^ERT2^ (WT) mice and Lgr5-GFP-Cre^ERT2^/Setdb1^lox/lox^ (Setdb1-KO) mice. Note the higher number of counts above the baseline in Setdb1-KO cells. (E) Genome Browser profile of normalized ATAC-seq reads and peaks-linked-to-genes (black bars) at the genomic region encompassing the aberrantly activated Itln-1 gene in tamoxifen-treated Lgr5-GFP-Cre^ERT2^ mice (WT) and in Lgr5-GFP-Cre^ERT2^/ Setdb1^lox/lox^ mice (Setdb1-KO). Arrows indicate ATAC-seq peaks gained in Setdb1-deficient cells.

**Figure 5.**
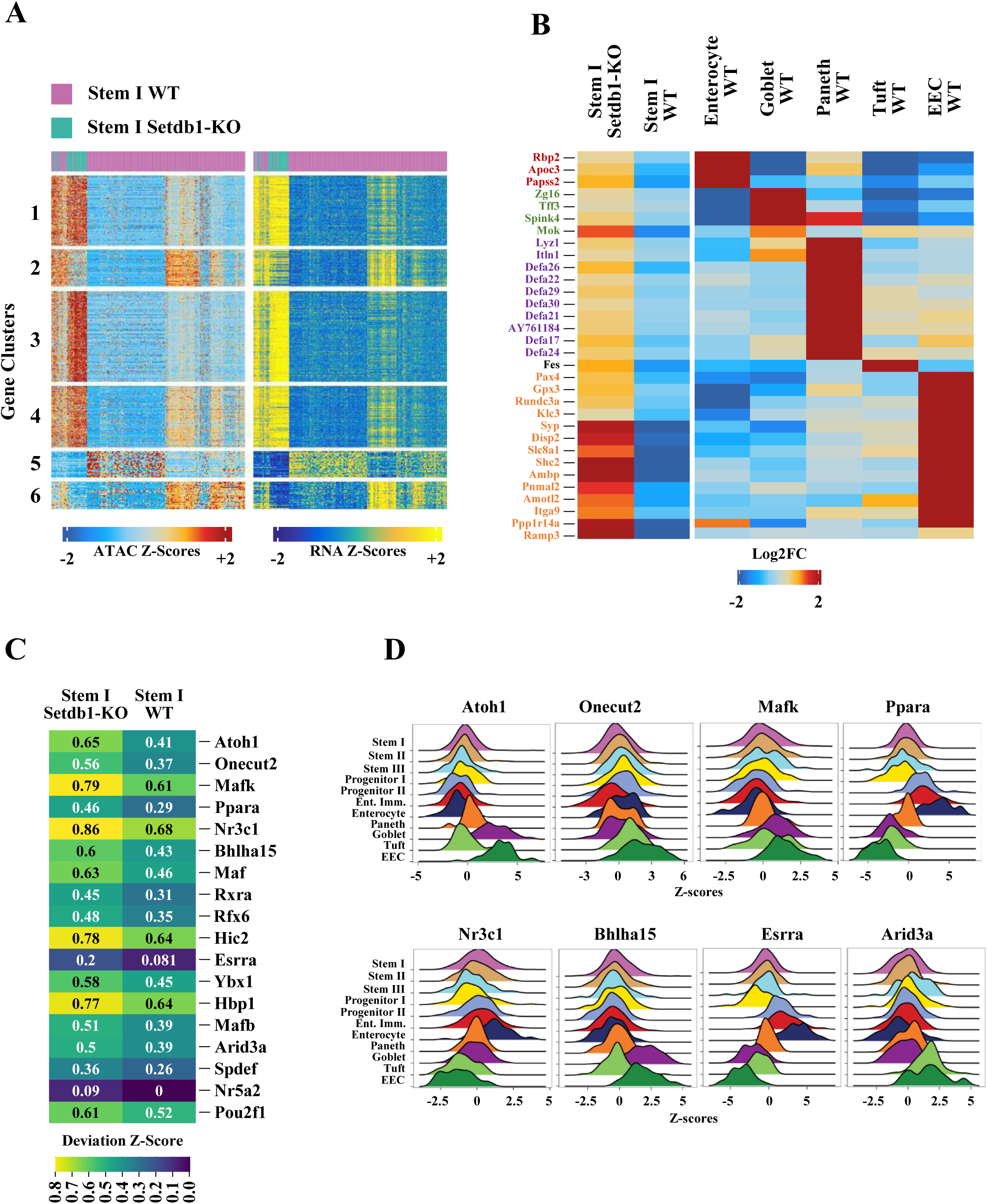
Aberrant activation of genes in Setdb1-deficient Lgr5^+^ stem cells. (A) Heatmaps and corresponding clustering of integrated RNA signals (panel at right) and normalized scATAC-seq signals (left panel) of the 4890 ATAC-seq peaks linked to 3634 unique genes in Stem-I cell population from tamoxifen-treated Lgr5-GFP-Cre^ERT2^ (WT; purple) and Lgr5-GFP-Cre^ERT2^/Setdb1^lox/lox^ (Setdb1-KO; green) mice. The six gene clusters are separated by space. The distribution of the total ATAC-seq peaks and linked to unique genes between the clusters is as follows: Cluster 1: 1097 peaks/815 genes; Cluster 2: 578 peaks/460 genes; Cluster 3: 1415 peaks/1001 genes; Cluster 4: 963 peaks/748 genes; Cluster 5: 409 peaks/313 genes; Cluster 6: 428 peaks/297 genes. Note the strong correlation between ATAC-seq signal and RNA levels in all of the gene clusters and the Setdb1-KO-specific increased chromatin accessibility and RNA expression in gene clusters 1, 3 and 4. Cluster 2 and 6 displays a mixed pattern of increased accessibility and RNA expression, while cluster 5 genes which are open and active in about 50% of wild type cells are less accessible and have decreased RNA levels in Setdb1-KO cells. (B) Gene expression intensity heatmap of aberrantly activated genes in Stem-I cell population from tamoxifen-treated Lgr5-GFP-Cre^ERT2^ (WT) and Lgr5-GFP-Cre^ERT2^/Setdb1^lox/lox^ (Setdb1-KO) mice. Panel at right shows the normalized intensity of the genes in in other epithelial cell clusters from wild-type mice. (C) Heatmap with normalized motif deviation Z-scores of Transcription Factors in Stem-I cell population of tamoxifen-treated Lgr5-GFP-Cre^ERT2^ (WT) and Lgr5-GFP-Cre^ERT2^/Setdb1^lox/lox^ (Setdb1-KO) mice. Transcription Factors whose binding motifs within the ATAC-seq peaks show significantly increased z-score values in Setdb1-deficient cells are listed. (D) Ridge plot of Transcription Factor motif accessibility z-scores across intestinal epithelial cell clusters. The density distribution of motif accessibility z-score for Nr1h3, Esrra, Atoh1, Nfya, Arid3a, Mafb, Bhlha15 and Phf21 is shown.

To gain mechanistic insights into how the above genes were activated prematurely in Stem-I cells, we performed Transcription Factor Motif enrichment analysis in the scATAC-seq regions detected in wild type and Setdb1-deficient Stem-I cells. We analyzed the enrichment of motifs beyond what is expected by chance, to reveal active binding by specific Transcription Factors at the accessible chromatin sites of interest. To correlate the motifs with the expression of the corresponding Transcription Factor, which was inferred via gene activity scores, we calculated deviation Z-scores and identified the positively correlating Transcription Factors, demonstrating a direct link between Transcription Factor abundance and regulatory activity in specific cell states (Figure 5C). We found that the *Setdb1* inactivation increased the motif accessibility and expression of 18 Transcription Factors in Setdb1-deficient Stem-I cells, which are normally enriched in one or more differentiated cell type. As shown in the Ridge plots of Figure 5D, Atoh1, Mafk and Bhlha15 TF motif exposure and expression is enriched in Goblet and EEC cells, Onecut2 motif in EEC cells, Ppara, Nr31c and Esrra in enterocytes and Arid3a in Tuft and EEC cells.

The above data suggest that premature exposure of TF binding sites in Setdb1-deficient Stem-I cells is sufficient to promote stochastic binding of the Transcription Factors, even if they are expressed at low levels. This provides a mechanistic basis for the observed premature activation of differentiated cell type-specific genes, which could affect differential gene expression and trajectory establishment. Such changes may result in increased transcriptome variations. We therefore tested cell-to-cell variability of gene expression patterns in all stem cell and Progenitor cell clusters. Consistent with the increased global heterogeneity of the scATAC-seq peak areas in Setdb1-KO cells, we observed significant changes of Coefficient Variations per Genes in cells within individual stem cell and Progenitor cell clusters, indicating that *Setdb1* inactivation results in significantly enhanced cell-to-cell variations of cellular transcriptomes (Figure 6A and 6B).

**Figure 6.**
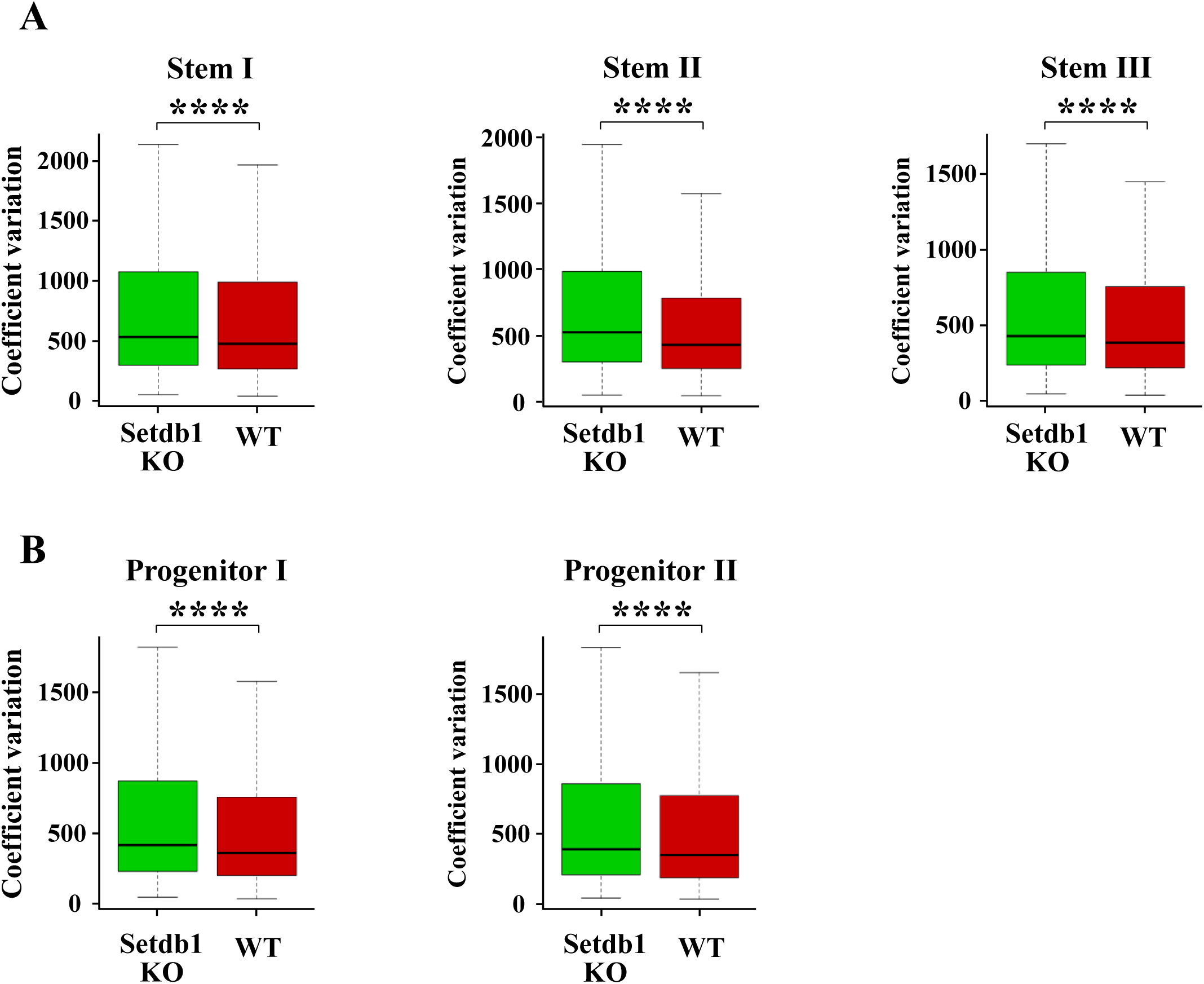
Increased cell-to-cell transcriptional variability Setdb1-deficient Lgr5^+^ Stem and Progenitor cell clusters. (A and B) Box plots showing Coefficient Variations per Gene in cells within Stem-I, Stem-II and Stem-III clusters (A) and within Progenitor-I and Progenitor-II clusters (B) from tamoxifen-treated Lgr5-GFP-Cre^ERT2^ (WT) and Lgr5-GFP-Cre^ERT2^/Setdb1^lox/lox^ (Setdb1-KO) mice. Mann-Whitney-Wilcoxon test, **** p < 2.2×10^-16^.

### Moderate reduction of H3K9 methylation inside and outside large heterochromatin domains correlate with the dynamic changes in the transposase-accessible chromatin areas in Setdb1-deficient cells

We compared the H3K9Me_]_profiles of FACS-sorted control and Setdb1-deficient Lgr5^+^-GFP cells. Consistent with the redundancy of Setdb1 function in generating H3K9 trimethylated nucleosomes^5,7,9,14^, we did not observe significant differences in H3K9Me_]_peaks between control and *Setdb1*-KO cells at the majority of large heterochromatin domains (Figure 7A, 7B, S6A and S6B).

**Figure 7.**
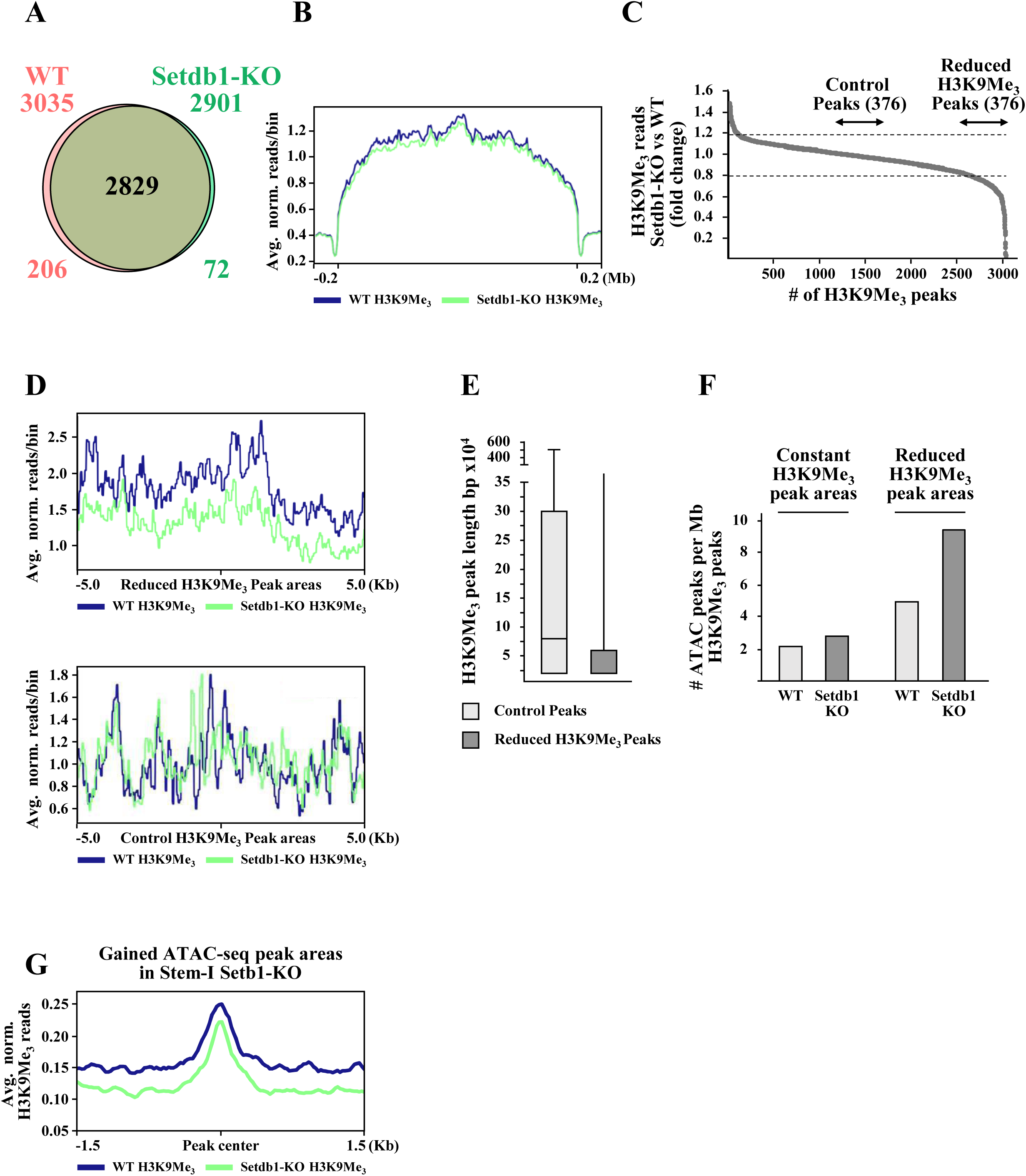
Differential effects of Setdb1 inactivation on H3K9Me_]_-containing long heterochromatin domains and shorter peak areas at the vicinity of transposase accessible regions (A) Venn diagram showing the extent of overlap of H3K9Me_]_CUT&Tag peaks between FACS-sorted GFP^+^ cells from tamoxifen-treated Lgr5-GFP-Cre^ERT2^ (WT) and Lgr5-GFP-Cre^ERT2^/Setdb1^lox/lox^ (Setdb1-KO) mice. The peaks were determined by SICER broad ChIP-seq peak caller tool. (B) Average distribution profiles of the CUT&Tag reads within the SICER-called H3K9Me_]_peaks indicated in (A). The graph shows average normalized reads per 50 bp bins. (C) Ranked coverage plots of the ratios of H3K9Me_]_reads within the SICER-called H3K9Me_]_peak areas between Setdb1-KO and WT cells. Dashed lines at +/-20% are indicated as base value of not significant change. The 376 Peaks below the 0.8-fold cutoff are called “Reduced H3K9Me_]_Peaks”. Control Peaks are randomly selected peaks with Setdb1-KO/WT H3K9Me_]_read ratios between 0.8 and 1.2-fold. (D) Average distribution profiles of H3K9Me_]_reads within the 376 selected H3K9Me_]_peaks containing reduced number of reads in Setdb1-KO cells (upper panel) and the 376 randomly selected control H3K9Me_]_peaks with unchanged H3K9Me_]_reads (bottom panel) indicated in (C). The graph shows average normalized reads per 50 bp bins. (E) Comparison of median peak lengths of the 376 control H3K9Me_]_peaks and those with reduced number of reads indicated in (C). (F) Evaluation of the overlaps between the peaks of the control and those with reduced H3K9Me_]_reads in (C), with ATAC-seq peaks in cells from tamoxifen-treated Lgr5-GFP-Cre^ERT2^ (WT) and Lgr5-GFP-Cre^ERT2^/Setdb1^lox/lox^ (Setdb1-KO) mice. (G) Comparison of the average distribution of H3K9Me_]_reads in the ATAC-seq peak areas gained in Setdb1-KO Stem-I cell cluster. Blue line shows H3K9Me_]_reads in FACS-sorted GFP^+^ cells from tamoxifen-treated Lgr5-GFP-Cre^ERT2^ (WT) mice. Greene line shows H3K9Me_]_reads detected in GFP^+^ cells isolated from tamoxifen-treated Lgr5-GFP-Cre^ERT2^/Setdb1^lox/lox^ (Setdb1-KO) mice. The graph shows average normalized reads per 50 bp bins.

However, comparing the H3K9Me_]_CUT&Tag reads within the individual called peaks, we detected a substantial (>20%) drop in 376 out of the total 3035 peaks (Figure 7C and 7D). Importantly, the length of these 376 H3K9Me_]_CUT&Tag peaks was significantly shorter and less variable, compared to the same number of randomly chosen peaks, in which H3K9Me_]_reads were not affected by *Setdb1* inactivation (Figure 7E).

Next, we compared the locations of the existing and the new ATAC-seq peaks in the Setdb1-deficient cells that overlap with H3K9Me_]_peaks. As expected, the vast majority of the ATAC-seq peak areas, corresponding to accessible open chromatin was excluded from the H3K9Me_]_peaks marking closed heterochromatin domains (Figure S6B). Interestingly however, the fraction of the ATAC-seq peaks that overlapped with the H3K9Me_]_peaks, was substantially enriched in the peak locations, with decreased H3K9Me_]_read content in Setdb1-deficient cells (Figure 7F). Next, we compared the H3K9Me_]_status in the newly formed ATAC-seq areas in *Setdb1*-KO Stem-I cells, using the H3K9Me_]_CUT&Tag data sets from GFP^+^ cells. Average coverage plots of H3K9Me_]_reads from GFP^+^ cells centered on the newly-formed ATAC-seq peak areas that were identified in Setdb1-deficient cells revealed substantial decrease of the H3K9Me_]_signals in Setdb1-deficient cells (Figure 7D).

Taken together, the above findings suggest that Setdb1-mediated H3K9 methylation regulates chromatin accessibility in a large number of gene-rich open genomic regions, while Setdb1 function in the majority of large heterochromatin regions is redundant.

## Discussion

Epigenetic enzymes regulate developmental programs through changes in chromatin structure, which influence the accessibility to transcriptional regulators.^7,11^ Regulation of chromatin accessibility involves dynamic events of heterochromatin compaction and structural transitions to open euchromatin, which control gene activity during ES cell differentiation to specific cell types or during developmental lineage specification.^9–13^

The main findings of this study demonstrate that the H3K9-methylase Setdb1, apart from its known function in the formation of large heterochromatin domains, plays an important role in controlling chromatin accessibility in open genomic regions in adult intestinal stem cells. This is demonstrated by the re-distribution of ATAC-seq peaks in Setdb1-deficient cells, which mark transposase accessible open chromatin domains. Our results suggest that mechanisms involving localized changes within established chromatin compartments are important for fine-tuning gene expression patterns, in addition to the known regulatory role of heterochromatin to euchromatin transitions and vice versa. Such delicate regulatory mechanisms may prevent spurious activation or repression of genes, thereby contributing to adult stem cell maintenance and proper differentiation. In line with this notion, we found that disruption of the tight control of chromatin accessibility in Lgr5^+^ intestinal stem cells leads to premature activation of genes whose expression is normally restricted to specific differentiated epithelial cell types. Premature gene activation is a consequence of premature exposure of transcription factor binding sites in gene regulatory regions, due to the loss of Setdb1-dependent localized chromatin modifications.

The most apparent consequence of the resulting aberrant gene expression profiles is increased cell-to-cell transcriptional variability within well-defined cellular clusters. Cell-to-cell heterogeneity in gene expression, discrete from the intrinsic noise inherently present in cell populations, has been recognized as a potentially important feature to explain biological phenomena.^36–38^ For example, it has been shown that increased cell-to-cell transcriptional variability is coupled with the aging-dependent response of CD4^+^ T cells to immune stimulation,^39^ in aging mouse muscle stem cells.^40^ Similar to the aging process, cellular differentiation is also tightly regulated by genetic and epigenetic factors and is modulated by stochastic signaling processes. Such multilevel control mechanisms generate temporally distinct transcriptomes, which characterize the identity of consecutive differentiation states. Since tight regulation is supposed to promote the creation of “more uniform” cellular transcriptomes, it is reasonable to assume that during the transition between the different cellular differentiation states, partial transcriptome diversification in individual cells may be part of the physiological differentiation process (schematically presented in Figure S7). Transition-specific variable transcriptomes must result from a stochastic equilibrium of regulatory factors and chromatin configurations, which determine the phenotypes of the previous and subsequent cellular states. Thus, we speculate that, temporary variations restricted to the transition periods may represent a hallmark feature of cellular differentiation and lineage specification.

Disruption of the control mechanisms, like those driven by the abnormal expansion of open chromatin domains in Setdb1-deficient cells, can halt the progression of differentiation, resulting in an increased number of cells trapped at the transition states. In agreement with this scenario, we observed abnormal, premature activation of a number of differentiation stage-specific genes in Setdb1-deficient Lgr5^+^ stem cells, which in turn elicit increased transcriptional variability in individual cells.

The consequences of abnormal gene activation may comprise alterations in stemness features and/or in the differentiation capacity of the cells. This possibility is supported by the strong dependence of Lgr5^+^ stem cells on Setdb1 function to generate intestinal organoid structures *in vitro*. The requirement of Setdb1 for the proper differentiation of intestinal stem cells, was further demonstrated by the findings of *in vivo* lineage tracing assays, where a dramatic accumulation of stem cell clusters was observed in mice carrying conditionally inactivated *Setdb1* alleles. The number of mature absorptive enterocytes was severely reduced in Setdb1-deficient Lgr5^+^ stem cell-derived progeny, in parallel with a comparable loss of cells in Progenitor II cell compartment. According to our trajectory analysis, this latter cluster appears to represent a unipotent transition state toward the enterocyte lineage. This suggests that in the absence of Setdb1 the above differentiation path is blocked at the very early stage during the transition of Stem cells to Progenitor II cells.

Interestingly, the secretory cell types in Setdb1-deficient Lgr5^+^ stem cell-derived progeny were readily detected, with marginal reduction of Paneth cells and marginal increase of Goblet cells, Tufts cells and Enteroendocrine cell types, with the exception of Goblet-I cell cluster, where a significant expansion was evident. Goblet-I cell cluster appears to be derived directly from Progenitor I/TA cells and corresponds to the immediate precursor of Goblet-II and Goblet-III clusters. The accumulation of both Progenitor-I/TA and Goblet-I cell types following *Setdb1* inactivation suggests that cells escaping the first block of the hierarchical differentiation paths are halted at a second differentiation checkpoint at the Progenitor cell state. Nonetheless, the existence of Lgr5^+^ stem-cell derived secretory and endocrine-related cell types in Setdb1-deficient mice suggests that these defects can be partially rescued by the activation of alternative lineage trajectories, through which they can arise directly from the Stem-I cell population. The activation of such alternative trajectories is likely to be facilitated by the increased transcriptional variability of cells that are blocked at intermediate differentiation checkpoints, which could provide cells with the flexibility to explore multiple differentiation directions (Figure S7).

Taken together, the results demonstrate that regulated chromatin accessibility impacts stage-specific transcriptome changes and plays an important role in executing proper differentiation paths in the intestinal epithelium. Perturbation of chromatin structure leads to aberrant and premature activation of genes. This results in increased cell-to-cell transcriptome variability for an extended period of time and in the inhibition of stem cell differentiation. Disruption of chromatin-controlled processes completely blocks differentiation into absorptive enterocytes. However, the activation of alternative differentiation paths compensates for the loss of secretory and endocrine-related cells, highlighting the remarkable plasticity in intestinal lineage specification regulation.

## RESOURCE AVAILABILITY

### Lead contact

Further information and requests for resources and reagents should be directed to and will be fulfilled by the lead contact, Iannis Talianidis

### Materials availability

Research materials generated in this study will be provided by the lead contact upon request.

### Data availability

Source Data files for the Total RNA-seq, scRNA-seq, scATAC-seq and CUT&Tag sequence data have been deposited in the Gene Expression Omnibus (GEO) under accession number GSE279796 as listed in the key resource table.

## ACKNOWLEDGEMENTS

We thank M. Koukaki and E. Moltsanidou for technical assistance and discussions during the course of the work; E. Stratidaki, and N. Gounalaki of the IMBB-Genomics facility and I. de la Rosa of the Core Genomics Unit of HMGU for assistance with the library preparations. This work was supported by the “Basic research Financing (Horizontal support of all Sciences)” under the National Recovery and Resilience Plan “Greece 2.0” funded by the European Union–NextGenerationEU, H.F.R.I. Project:15276 (to IT and M.D.L), the AXA Research Fund Chair in Epigenetics Program #2016 (to IT); the CIDEGENT program: CIDEXG/2023/30 and internal funds from the Helmholtz Pioneer Campus (to C.P.M-J).

## AUTHOR CONTRIBUTIONS

I.P. and I.T. conceived and designed the study. I.P., I.K.D. and H.K. performed experiments and evaluated the results. D.B., M.S. and G.G. performed bioinformatics analyses of the data. E.D. analyzed the data. C.P.M-J, M.D.L. and I.T. supervised the project. I.T. wrote the manuscript. All of the authors reviewed and edited the manuscript.

## DECLARATION OF INTERESTS

The authors declare no competing interest.

## STAR METHODS

### Key resources table

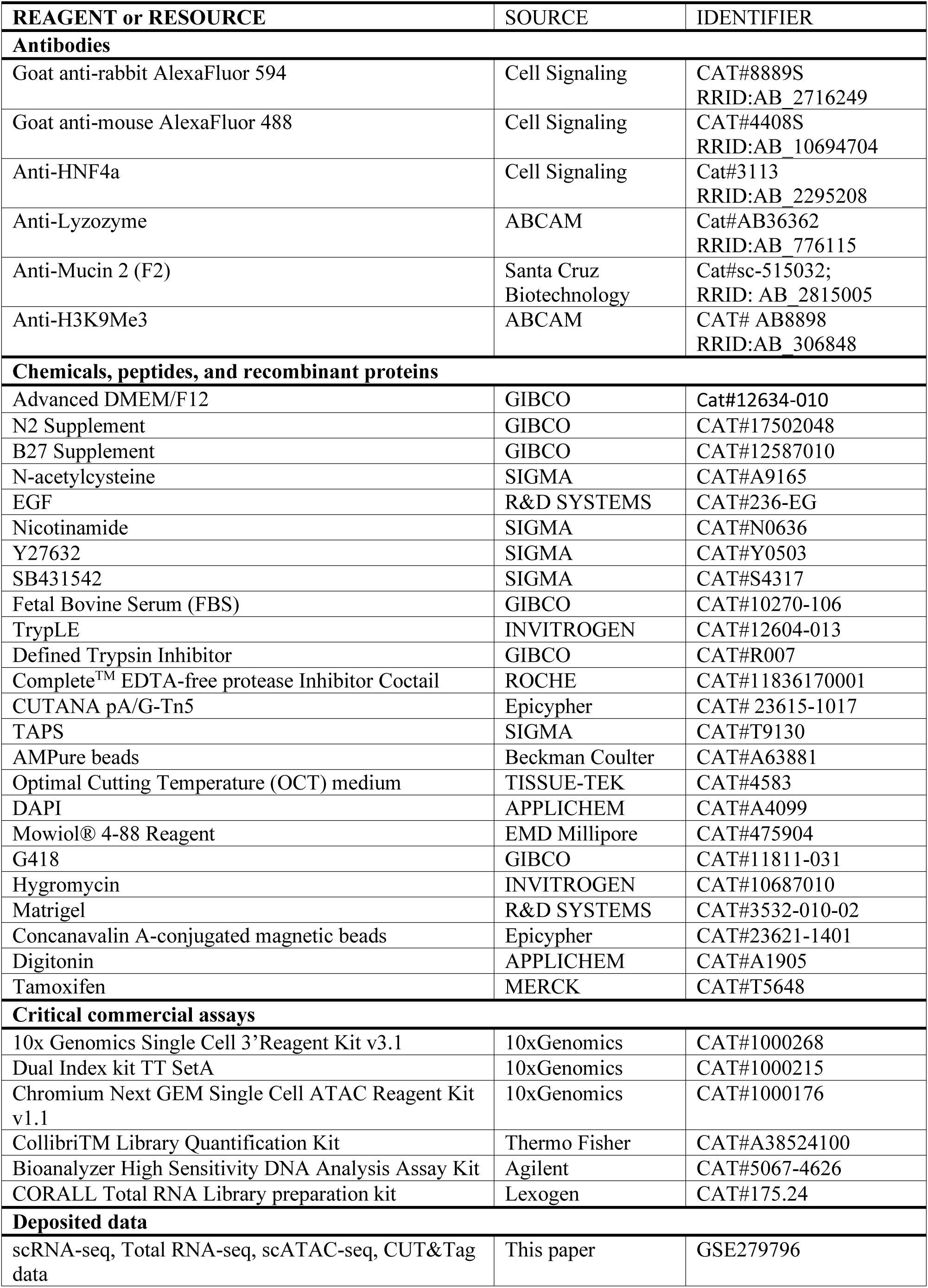
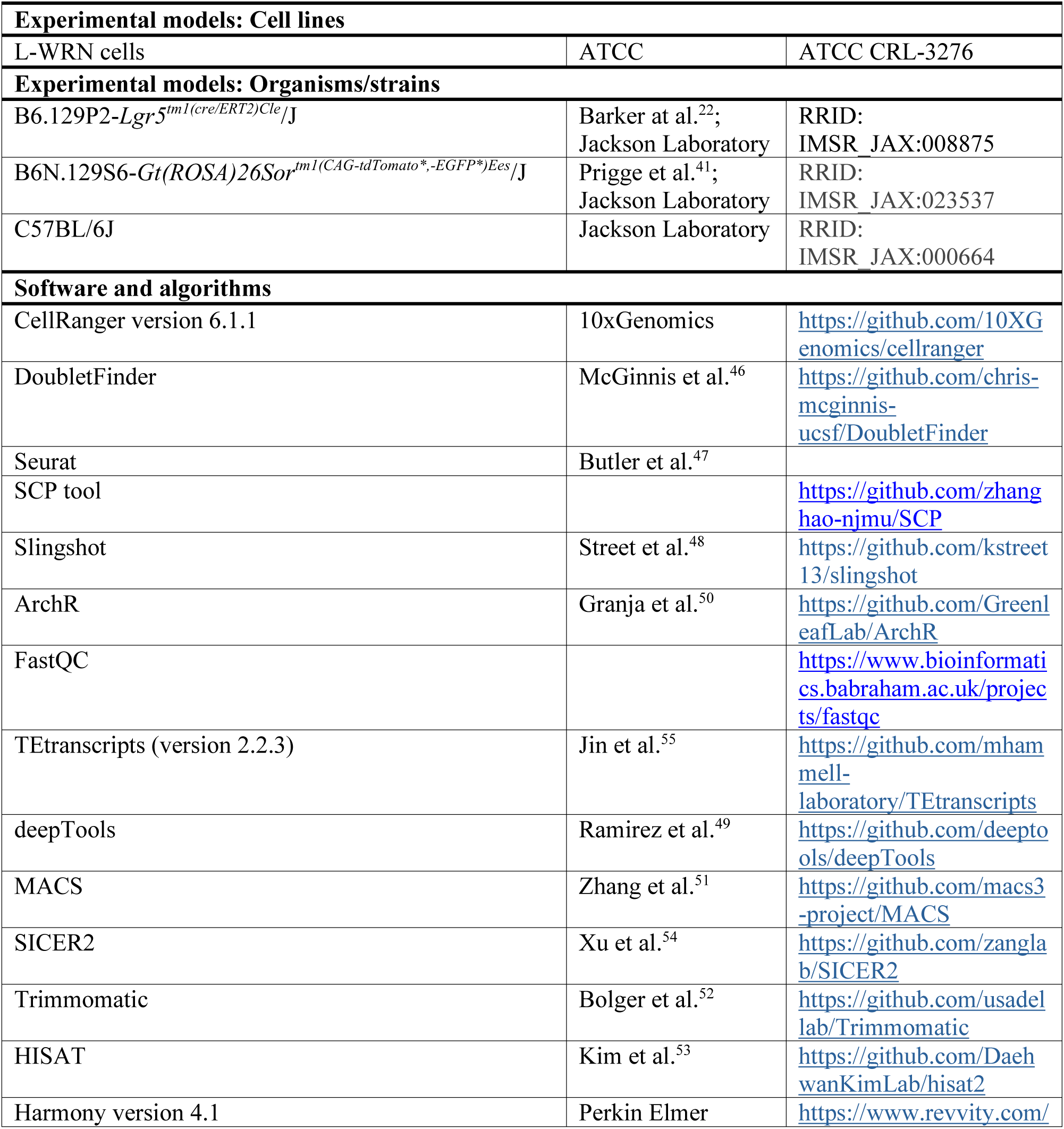

## EXPERIMENTAL MODEL AND STUDY PARTICIPANT DETAILS

### Mice

Lgr5-GFP-Cre^ERT2^ (B6.129P2-*Lgr5^tm1(cre/ERT2)Cle^*/J)^22^ and ROSA(CAG-tdTomato*,EGFP*)Ees (called nTnG) (B6N.129S6-*Gt(ROSA)26Sor^tm1(CAG-tdTomato*,-EGFP*)Ees^*/J)^41^ mice were obtained from Jackson Laboratory. Setdb1^lox/lox^ mice carrying floxed exon 3 alleles were generated by crossing Setdb1^tm1a(EUCOMM)Wtsi^ mice, obtained from Wellcome Trust Sanger Institute, with B6.129S4-*Gt(ROSA)26Sor^tm2(FLP*)Sor^*/J mice, obtained from Jackson Laboratory. After verifying flp recombinase-mediated deletion and recombination of the FRT site-flanked LacZ/neomycin cassette, the flp transgene was removed by serial back-crossing with C57Bl6 mice. Lgr5-GFP-Cre^ERT2^/ Setdb1^lox/lox^, Lgr5-GFP-Cre^ERT2^/nTnG and Lgr5-GFP-Cre^ERT2^/ Setdb1^lox/lox^/nTnG mice were obtained by crossing the respective mouse strains, as described in Figure S1B, S1C and S1D, respectively. The resulting mice were fertile and viable over 8 months of age. Mice were kept in grouped cages in a temperature-controlled, pathogen-free facility on a 12-hour light/dark cycle and fed with standard chow diet containing 19% protein and 5% fat (Altromin 1324) and water ad libitum. All animal experiments were approved by the Ethical Review Board of IMBB-FORTH and the Animal Ethics Committee of the Prefecture of Crete and were performed in accordance with the respective national and European Union regulations. All experiments were performed in randomly chosen age-matched male mice. No blinding was used in this study. Unless otherwise indicated, mice were treated and analyzed at 60-65 days after birth. Treatment of mice were performed with intraperitoneal injections of either vehicle (corn-oil) or 70 mg/kg Tamoxifen.

## METHOD DETAILS

### Isolation of intestinal epithelial cells

To isolate crypts, the ileum was dissected and washed with cold phosphate buffered saline (PBS) using blunt-ended syringe, opened longitudinally with a scissor and cut into 3-5 mm pieces. After extensive washing with cold PBS, the tissue fragments were resuspended in 30 volumes of cold PBS, containing 20 mM EDTA and incubated with constant shaking for 30 min at 4^0^C. The tissue fragments allowed to settle by gravity to remove supernatant and resuspended in cold PBS. After vigorous shaking and trituration with 10ml pipette the tissue pieces were collected by gravity sedimentation. The supernatant was discarded and the PBS extraction was repeated 7 to 9 times, until the supernatant contained mainly crypts, as verified by microscopic examination. The final supernatant containing enriched crypt fraction was passed through 70 μm cell strainer and centrifuged at 400g for 5 min. To isolate epithelial cells, the crypt pellets were incubated in 10 volumes of preheated TrypLE Express (Gibco) for 4.5 min at 37 ^0^C. The reactions were stopped by the addition of equal volume of Defined Trypsin Inhibitor (Gibco). The cells were passed through 40 μm cell strainer, centrifuged at 400g for 5 min and resuspended in ice-cold PBS containing 1 mM EDTA and 2% FBS. GFP^+^ cells were sorted using BD FACSAria III cell sorter gated by forward scatter, side scatter and pulse width parameter at a flow rate of 3-5000 events/second.

### Immunostaining assays

For immunofluorescence staining, freshly isolated ileal fragments were cut longitudinally and washed with PBS. The tissue fragments were prefixed in 4% paraformaldehyde (PFA) in PBS for 2 hours at room temperature and treated 30% ice cold sucrose, before embedding in Optimal Cutting Temperature (OCT) embedding medium and were frozen in liquid nitrogen. Frozen sections (5 to 7 μm thick) were air dried blocked in 5% BSA in PBS containing 0.1% Triton X-100 for 1 hour and then incubated with 1% BSA, 0.1% Triton X-100 in PBS with the indicated primary antibodies at 4^0^C overnight. After incubation with AlexaFluor 594-or AlexaFluor 488-conjugated goat anti-rabbit or anti-mouse secondary antibodies for 1 hour at room temperature and counterstained with 1 μg/ml DAPI for 10 min, the slides were covered with Mowiol® 4-88 Reagent (EMD Millipore, 475904), as described.^43^ Fluorescence images were observed using a Leica TCS SP8 confocal microscope.

### High Content Microscopy analyses of wide field and confocal images

Isolated intestinal epithelial cells were plated onto glass cover slips and fixed with 4% paraformaldehyde (PFA). Following antibody staining, immunofluorescence images were acquired with a 20x or 40x lens (Olympus, Shinjuku City, Tokyo, Japan) with an Operetta High Content Screening Microscope (HCSM, PerkinElmer, Waltham, MA, USA) in wide field and analyzed using Harmony software v4.1.

To estimate relative protein expression levels, antibody-stained epithelial cells were segmented based on DAPI nuclear staining. Clusters of adjacent cells were excluded from the analysis. Among the remaining cells, only GFP-positive populations were selected, and for these, the nuclear sum intensity of the target protein signal was quantified. The resulting values were plotted using GraphPad Prism v6.

### Organoid cultures

About 500 FACS-sorted GFP^+^ cells were resuspended in 50 μl Matrigel and applied to pre-warmed tissue culture plates to form hemispherical droplets. Following solidification, medium containing Advanced DMEM/F12, 2 mM Glutamax, 10 mM Hepes, 100 U/ml penicillin, 100 μg/ml streptomycin, N2 Supplement (1x), B27 Supplement (1x), 1 mM N-acetylcysteine, 50 ng/ml EGF, 10 mM Nicotinamide, 10 μM Y27632, 10 μM SB431542 and 50% conditioned WRN medium was added to cover the Matrigel domes. The medium was changed every 2 days for the indicated times. Conditioned WRN medium was prepared from L-WRN (ATCC CRL-3276) cells expressing Wnt3A, R-spondin and Noggin^42^. The cells were grown in DMEM/10%FBS supplemented with 0.5 mg/ml G418 and 0.5 mg/ml Hygromycin. To prepare conditioned medium, the cells were split and grown in DMEM/10%FBS, without the selection antibiotics until confluence. Fresh DMEM was added without FBS and the cells were incubated for 24 hours at 37^0^C. The supernatant was collected and the plates were replaced with new medium. Supernatants from 4 successive 24-hours incubations were collected and stored in-20 ^0^C. For passaging organoids, the cells in Matrigel were resuspended in 500 μl TrypLE Express supplemented with 10 μM Y-27632 and incubated for 5 min at 37 ^0^C. After the addition of equal volume of Defined Trypsin Inhibitor, the suspension was extensively triturated using glass Pasteur pipette. After counting the cells were re-seeded using Matrigel.

### CUT&Tag assays

CUT&Tag assays were performed as described previously.^44^ Briefly, nuclei were prepared from the isolated, FACS-sorted GFP^+^ intestinal epithelial cells by resuspension in 10 volumes of ice-cold Swelling Buffer, containing 10 mM Tris pH 7.5, 2 mM MgCl_]_, 3 mM CaCl_]_and protease inhibitor cocktail (Complete^TM^ EDTA-free, Roche). After centrifugation at 400g for 5 min at 4^0^C, the pellet was resuspended in Lysis Buffer, containing 10 mM Tris pH 7.5, 2 mM MgCl_]_, 3 mM CaCl_]_, 10% glycerol, 1% NP40 and protease inhibitor cocktail. After incubation in ice for 3 min the nuclei were collected by centrifugation for 5 min at 400g, at 4^0^C, washed twice with PBS. Then, 10^5^ nuclei were resuspended in a buffer containing 20 mM HEPES–KOH pH 7.9, 10 mM KCl, 0.1% Triton X-100, 20% Glycerol, 0.5 mM Spermidine, protease inhibitor cocktail and incubated for 10 min with Concanavalin A-conjugated magnetic beads (Epicypher) that were previously activated by repeated washings with a buffer containing 20 mM HEPES pH 7.9, 10 mM KCl, 1 mM CaCl_]_, 1 mM MnCl_]_. The reactions were supplemented with 0.5 volume of a buffer containing 20 mM HEPES-NaOH, pH 7.5, 150 mM NaCl, 0.5 mM Spermidine, protease inhibitor cocktail, 0.01% Digitonin, 2 mM EDTA and 0.5 μg of the anti-H3K9Me_]_or IgG negative control primary antibodies. The samples were incubated overnight at 4^0^C by gentle agitation. The beads were then supplemented with a buffer containing 20 mM HEPES-NaOH, pH 7.5, 150 mM NaCl, 0.5 mM Spermidine, protease inhibitor cocktail, 0.01% Digitonin and 0.5 μg of anti-rabbit secondary antibody (Epicypher). After incubation at room temperature for 30 min, the beads were washed twice with the same buffer and resuspended in the same buffer containing 300 mM NaCl. Protein A/G-fused Tn5 transposase protein (CUTANA pA/G-Tn5, Epicypher) was added and the samples were incubated for 1 hour at room temperature followed by washing with a buffer containing 20 mM HEPES-NaOH, pH 7.5, 300 mM NaCl, 0.5 mM Spermidine, protease inhibitor cocktail and 0.01% Digitonin. The beads were collected and resuspended in the above buffer containing 10 mM MgCl_]_, incubated for 1 hour at room temperature, washed once with 10 mM TAPS pH 8.5 and 0.2 mM EDTA and incubated with 10 mM TAPS pH 8.5 and 0.1% SDS for 1 hour at 58^0^C, to release DNA. After quenching by the addition of 3 volumes of 0.67% Triton-X100, the samples were subjected to PCR reaction (14 cycles) using universal i5 primers and barcoded i7 primers. After cleanup with AMPure beads (Beckman Coulter), quantification and quality evaluation in Agilent Bionalyzer, the libraries were sequenced using Illumina NextSeq500 platform.

### Single-cell RNA-seq sample preparation and sequencing

After isolation of GFP^+^ cells using BD FACSAria III cell sorter, viable cells were fixed using 10% DMSO in media and stored at-80^0^C until further use. Τwo independent biological replicates were used for Lgr5^+^WT and Lgr5^+^Setdb1KO mice respectively. Cells were washed twice in PBS/ BSA 0,02% and centrifuged at 300 × g for 5 min at 4^0^C. The samples were processed according to the 10x Genomics Single Cell 3′ Reagent Kit User Guide. cDNA was assessed using a Bioanalyzer High Sensitivity DNA Analysis assay (Agilent) and the libraries were quantified using the Collibri™ Library Quantification Kit (Thermo Fischer Scientific, A38524500) in a QuantStudio™ 6 Flex Real-Time PCR System (Thermo Fisher Scientific). The pooled library was sequenced using SP100 flow cells in a NovaSeq6000 sequencer (Illumina) at the HMGU Core Facility for NGS Sequencing at a sequencing depth of 40,000 reads per cell.

### Single-cell ATAC-seq library preparation and sequencing

A single-cell paired-matched analysis of the assay for transposase-accessible chromatin with sequencing (scATAC-seq) was performed with approximately 300,000 pre-sorted cells from the same cell suspension that were used for scRNAseq library preparation. The cells were pelleted by centrifugation at 300 × g for 5 min at 4^0^C, washed twice with pre-chilled 1 ml of PBS/0.2%BSA and resuspended in 100 ml of Lysis buffer for 5 minutes, according to the Nuclei Isolation for Single Cell ATAC Sequencing demonstrated protocol (CG000169, 10X Genomics). The isolated nuclei were counted under a microscope and proceeded to Tagmentation reaction according to the manufacturer’s protocol (Chromium Next GEM Single Cell ATAC Reagent Kits v1.1, manual CG000209, RevD). Briefly the nuclei after transposition were loaded in a chip H and after GEM generation and clean-up, they were subjected to 9 cycles indexing PCR reaction. The final scATAC libraries were assessed using a Bioanalyzer High Sensitivity DNA Analysis assay (Agilent) and the Collibri™ Library Quantification Kit as above. Sequencing was performed in a SP 100 flow cell in a NovaSeq 6000 sequencer (Illumina) at the HMGU Core Facility for NGS Sequencing with the sequencing length recommended by 10x Genomics (50, 8, 16, 50). On average, 50,000 reads per nucleus were yielded.

### Single-Cell RNA-seq Data Analysis

Initial preprocessing was performed using the CellRanger pipeline (version 6.1.1, 10x Genomics), including alignment to the reference genome, barcode demultiplexing, and unique molecular identifier (UMI) counting. The CellRanger web summary report provided metrics such as the estimated number of cells, median genes per cell, and the percentage of reads mapping to the genome^45^.

To identify and remove potential doublets, the DoubletFinder algorithm^46^ was applied using the following parameters: pN = 0.25, pK = 0.09, the expected number of doublets (nExp), and the first 10 principal components (PCs = 1:10). The doublet detection rate was based on nExp, calculated from the estimated number of cells and the experimental protocol. Cells identified as doublets were excluded from further analysis.

Post-doublet removal, quality control filtering was performed using the Seurat package (version 5.0.3)^47^ to retain high-quality cells. Cells were filtered based on the following criteria: a number of detected features (genes) per cell between 200 and 4000, a mitochondrial gene expression percentage below 5%, and a total RNA count not exceeding 18,000.

Normalization was conducted using the “LogNormalize” method, followed by the identification of the top 2,000 highly variable genes with the variance-stabilizing transformation (vst) method. The data were then centered and scaled, and principal component analysis (PCA) was used for dimensionality reduction. A nearest neighbor graph was constructed using the top 10 principal components, and cells were clustered using the Louvain algorithm at a resolution of 0.5 via the Seurat FindClusters function. Clustering was visualized with Uniform Manifold Approximation and Projection (UMAP). The resulting Seurat object was then input into the SCP (Single-Cell Pipeline) tool (https://github.com/zhanghao-njmu/SCP) for visualization and trajectory inference using the Slingshot algorithm.^48^

### Assessment of cell-to-cell transcriptome variation

Cell-to-cell heterogeneity in mRNA expression was assessed as described before^39^ by calculating the coefficient of variation (CV) in gene expression separately for wild-type and knockout cells. Gene count matrices were generated for each condition, and the CV for each gene was computed as the standard deviation of gene expression divided by the mean expression, multiplied by 100. Boxplots were generated to visualize differences in CV between conditions, and statistical significance was assessed using a Wilcoxon rank-sum test.

### Single-cell ATAC-seq data analysis

Initial preprocessing was performed using the CellRanger pipeline (version 6.1.1, 10x Genomics), including alignment of reads to the reference genome and generation of BAM files. The BAM files were normalized using the *bamCoverage* tool from deepTools (version 3.3.2)^49^ with the BPM (Bins Per Million) method. We filtered out cells with less than 2.000 fragments and cells with TSS enrichment less than 4. For dimensionality reduction, Latent Semantic Indexing (LSI) was employed using the *addIterativeLSI* function of ArchR.^50^ Cells were clustered based on the reduced LSI space using a k-nearest neighbors (kNN) algorithm (*addClusters* function). The resolution of clustering was determined empirically to maximize biological relevance. Gene scores were calculated with the *addGeneScoreMatrix* function, which estimates gene expression by computing accessibility of regulatory elements near gene bodies and promoters. Integration of scATAC-seq and scRNA-seq data was carried out using ArchR’s *addGeneIntegrationMatrix* function. Gene expression profiles from matched scRNA-seq data were mapped to scATAC-seq cells by comparing the gene score matrix from scATAC-seq with the gene expression matrix from scRNA-seq. Cells were paired based on similarity in their profiles, allowing the assignment of gene expression information to the scATAC-seq cells.

Peak calling of scATAC-seq was performed as follows. After the integration was made, we performed peak calling per cluster per condition. Using the functions of *addGroupCoverages* and *addReproduciblePeakSet* ArchR first builds a coverage profile for each group of cells and then calls Peaks using MACS2.^51^ Then, with the *getBW* function and *ReadsInTSS* normalization method bigwig files were generated that were used for the cluster specific browser plots (Figure 4E) and the intensity heatmaps (Figure 4A). Intensity plots of ATAC-seq signals were generated using the *computeMatrix* function of *deeptools* with reference points. To create the reference peak list, we used the union of common and unique peaks of Stem-I-WT and Stem-I-KO cells obtained by the function intersect and subtract of bedtools (Figure 4A).

### Peak-to-Gene Linkage Analysis

To explore potential regulatory relationships between chromatin accessibility and gene expression, peak-to-gene linkage analysis was performed using the *addPeak2GeneLinks* function in ArchR^50^ with a maximum distance of 250.000bp, a correlation cutoff equal to 0.45 for only the Stem-I-KO and Stem-I-WT cells. This analysis identifies correlations between peaks and gene expression by considering accessibility and expression values across cells. The fact that two elements are only accessible in a particular cell community does not necessarily mean that they have a regulatory relationship with each other. However, when a reachable peak is continuously accessible whilst a nearby gene is transcribed, we can deduce that this peak is a potential regulatory transcriptional element of the associated (linked) gene. Finally, results were visualized with the *plotPeak2GeneHeatmap* function, which displays the peak-to-gene links for cells within the Stem-I cluster using an FDR cutoff of 1e-04 (Figure 5A). When we used the *addPeak2GeneLinks* function for only the Stem-I-WT and then only for the Stem-I-KO we could generate the number of peaks that were associated for each gene in these two conditions. Cumulative distribution analysis (R) allows to represent the proportion of regions with more or less than a specific threshold value for peak-to-genes observations (number of ATAC regions associated with one given gene).

### Motif analysis and identification of transcriptional regulators

Motif annotations of our scATAC-seq regions were added using the **CIS-BP** motif database (*addMotifAnnotations*, and background peak sets were generated with *addBgdPeaks* to enable bias-corrected analyses. ChromVAR deviation scores were calculated with *addDeviationsMatrix* to quantify motif accessibility deviations across groups. Correlation analyses between motif accessibility (MotifMatrix) and gene expression (GeneIntegrationMatrix) were performed with *correlateMatrices*. Motifs were classified as active transcription factors if they displayed strong positive correlation with gene expression (correlation > 0.5), statistical significance (p adjusted < 0.05). Motif activity z-scores of active Transcription Factors were aggregated by cell type and condition using getGroupSE and were plotted normalized (Figure 5C) and as a distribution (Figure 5D).

### Analysis of H3K9me_]_CUT&Tag data

For the analysis of H3K9me_]_CUT&Tag data, the quality of all FASTQ files was assessed using FastQC (https://www.bioinformatics.babraham.ac.uk/projects/fastqc). Low-quality bases and adapter sequences were trimmed using Trimmomatic (version 0.39)^52^. The cleaned FASTQ files were aligned to the UCSC mm10 mouse genome using HISAT2 (version 2.1.0)^53^ with default settings, retaining only reads with a mapping quality score greater than 30. BAM files were converted to BigWig format using deepTools (version 3.3.2)^49^ and visualized in the UCSC Genome Browser. Read counts were standardized by down sampling to match the sample with the fewest reads, and peak regions were identified using SICER2.^54^

Average coverage profiles across peak centers of H3K9me_]_and ATAC-seq data were computed and visualized using DeepTools.^49^ The computeMatrix function was used to calculate the signal across genomic regions, with a bin size of 50 bp. The matrix was generated from normalized BigWig files, providing coverage data at each bin across specified regions. The resulting signal profiles were then visualized using the plotProfile function to represent the average signal distribution.

### Total RNA purification and RNA-sequencing for retroviral transcript detection

Total RNAs were purified from the FACS-sorted GFP^+^ epithelial cells using Trizol extraction, as described previously.^43^ Briefly, the cells were resuspended in 10 volumes of Trizol reagent by brief vortexing, followed by incubation at room temperature for 5 min. After the addition of 0.2 volumes of chloroform and further incubation at room temperature for 3 minutes, the samples were centrifuged at 12000 g for 15 minutes at 4^0^C and the aqueous phase was collected and precipitated by the addition of equal volume of isopropanol. After 10 minutes’ incubation at room temperature, the RNA was collected by centrifugation at 12000g for 15 minutes. The pellet was resuspended in H_]_O and re-precipitated with ethanol. The RNA samples were further purified by digestion with 10 units of DNase-I for 10 min at 37^0^C, followed by purification with phenol/chloroform extraction and ethanol precipitation.

To obtain maximal 5’ coverage, RNA-seq libraries were generated using CORALL Total RNA Library preparation kit from Lexogen and sequenced in an Illumina NextSeq 500 system.

Transposable element (TE) and gene expression levels were quantified using TEtranscripts (version 2.2.3)^55^. RNA-seq data were aligned to the reference genome, and TEtranscripts was used to estimate the expression of both TEs and protein coding genes. Following the differential expression analysis, RPKM (Reads Per Kilobase of transcript, per Million mapped reads) values were calculated for comparisons of individual TE transcripts and protein coding transcripts.

### Quantification and statistical analysis

Comparisons between two groups were performed using Mann-Whitney U-test or Mann-Whitney-Wilcoxon test as indicated in the legends. scRNA-seq, scATAC-seq, CUT&Tag data were obtained from at least two biological replicates.

## Code availability

https://github.com/GCMLab-Forth/scRNA_scATAC_analysis

---

**Figure S1.**
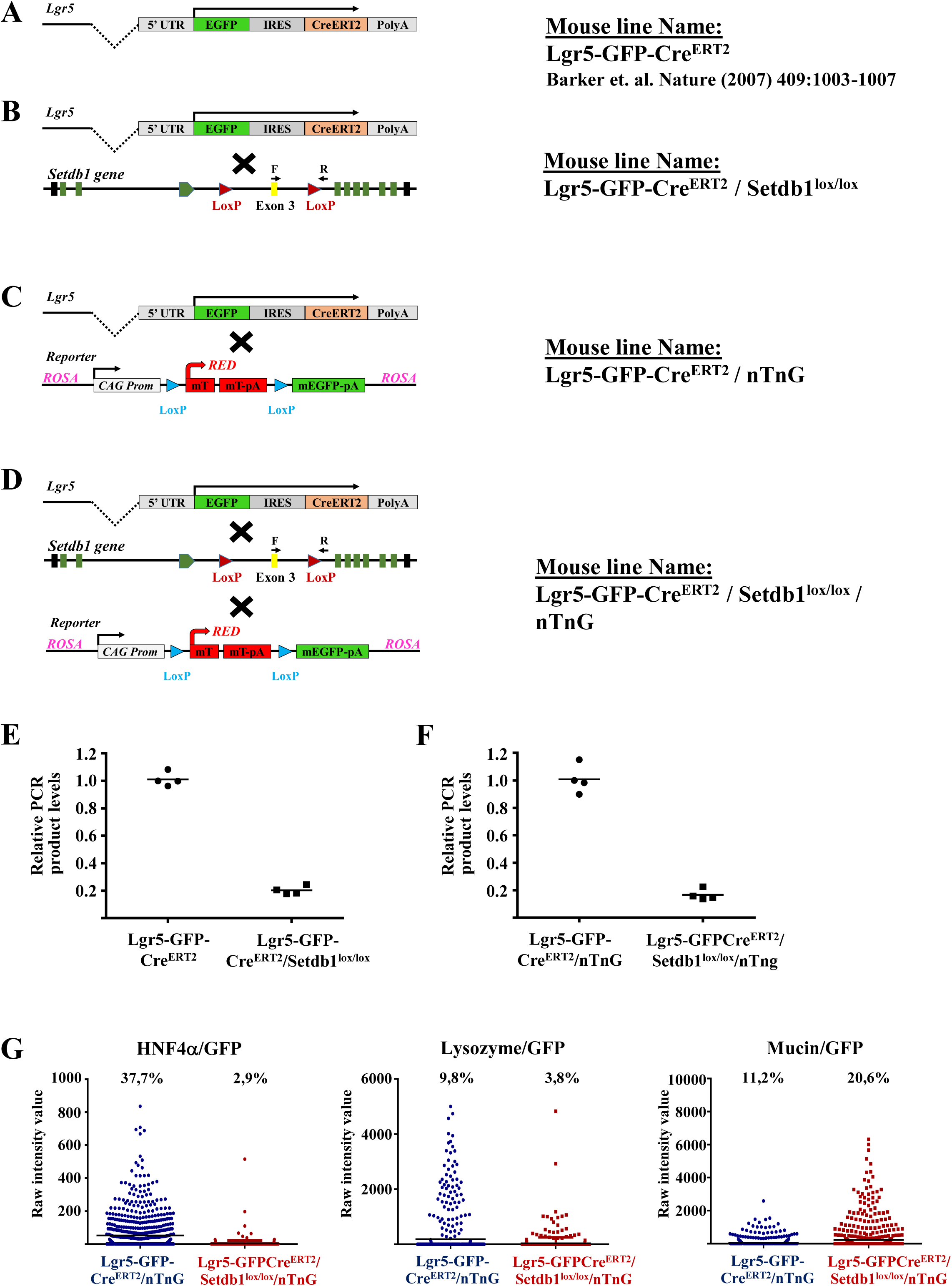
Schematic presentation of the genotypes of mouse models and analysis of the extent of the deletion of Setdb1-exon 3. (A) Lgr5-GFP-Cre^ERT2^ mice described in Ref 22, contains one knock-in allele expressing GFP and Cre^ERT2^ specifically in Lgr5^+^ stem cells. (B) Crossing Lgr5-GFP-Cre^ERT2^ mice with Setdb1^lox/lox^ mice generates Lgr5-GFP-Cre^ERT2^/ Setdb1^lox/lox^ mice, which upon tamoxifen treatment results in the deletion of exon 3 of Setdb1 and the generation of a premature stop codon in the area of exon 4, specifically in Lgr5^+^ stem cells. (C) Crossing Lgr5-GFP-Cre^ERT2^ mice with ROSA(CAG-tdTomato*EGFP*)Ees mice generates mice named Lgr5-GFP-Cre^ERT2^/nTnG, which express tomato-red in all cells and GFP and Cre^ERT2^ specifically in Lgr5^+^ stem cells. Upon tamoxifen treatment, all Lgr5^+^ stem cell progeny will express GFP from both Lgr5 and the ROSA locus. In parallel, these cells also loose tomato-red expression. (D) Crossing Lgr5-GFP-Cre^ERT2^ mice with Setdb1^lox/lox^ and ROSA(CAG-tdTomato*EGFP*)Ees mice, generates mice named Lgr5-GFP-Cre^ERT2^/ Setdb1^lox/lox^/nTnG. These mice express tomato-red in all cells and GFP and Cre^ERT2^ specifically in Lgr5^+^ stem cells. Upon tamoxifen treatment, exon 3 of Setdb1 is deleted specifically in Lgr5^+^ stem cells and all Setdb1-deficient Lgr5^+^ stem cell progeny will express GFP from both Lgr5 and the ROSA locus. (E-F) Deletion of Setdb1 exon 3 was verified by PCR from FACS-sorted GFP^+^ cells of Tamoxifen-treated Lgr5-GFP-Cre^ERT2^ and Lgr5-GFP-Cre^ERT2^/ Setdb1^lox/lox^ (E) or Lgr5-GFP-Cre^ERT2^/nTnG and Lgr5-GFP-Cre^ERT2^/ Setdb1^lox/lox^/nTnG (F) mice, using a primer pair hybridizing to sequences inside Exon 3 (F) and outside the LoxP site (R). The data from 4 cell preparations and mean their values are presented as relative to the values obtained with cells from Lgr5-GFP-Cre^ERT2^ and Lgr5-GFP-Cre^ERT2^/nTnG mice. (G) Estimation of HNF4a+ enterocytes, Lysozyme+ Paneth cells and Mucin+ Goblet cells in GFP+ cell populations from Tamoxifen-treated Lgr5-GFP-Cre^ERT2^/nTnG and Lgr5-GFP-Cre^ERT2^/ Setdb1^lox/lox^/nTnG. Intestinal epithelial cells from the mice were isolated and plated onto glass coverslips. The coverslips were stained with antibodies against HNF4a, Lysozyme and Mucin2. The graphs show High content Microscopy (HCM) measurements of individual cells double stained with the above markers and with GFP antibody from 786 cells in HNF4a/GFP-stained samples, from 802 cells in Lysozyme/GFP samples and 819 cells in Mucin/GFP-stained samples. Only cells with sum-intensity above background and with proper intracellular localization were counted. Numbers indicate the percentage of cells with staining intensity above threshold.

**Figure S2.**
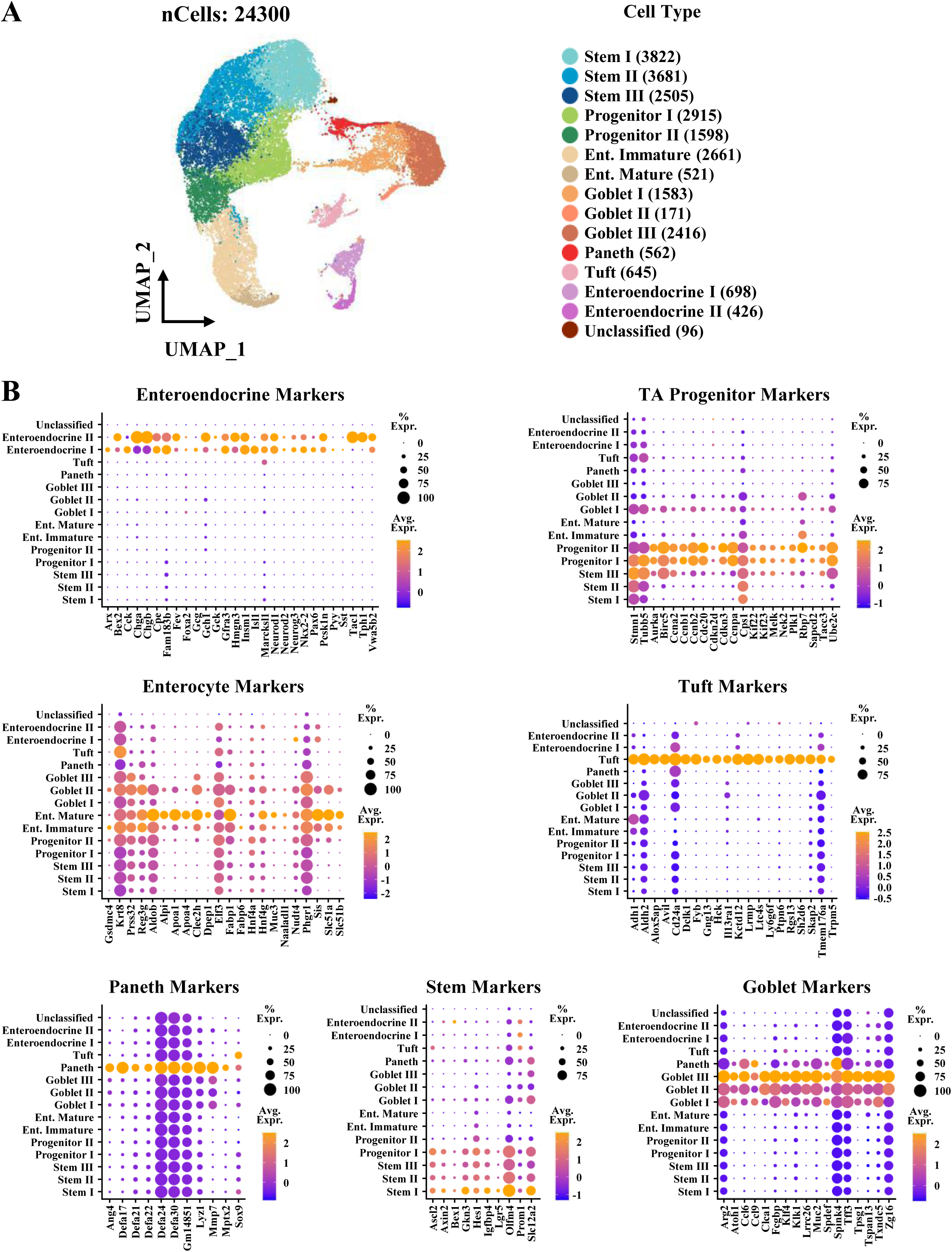
Identification of intestinal epithelial cell types. **a** UMAP projection of 24.300 single cells (merged datasets) identifies cell types based on the expression of known marker genes. The color codes of the annotated cell types are shown at the right. **b** Dot blot analysis of marker gene expression in the different cell clusters.

**Figure S3.**
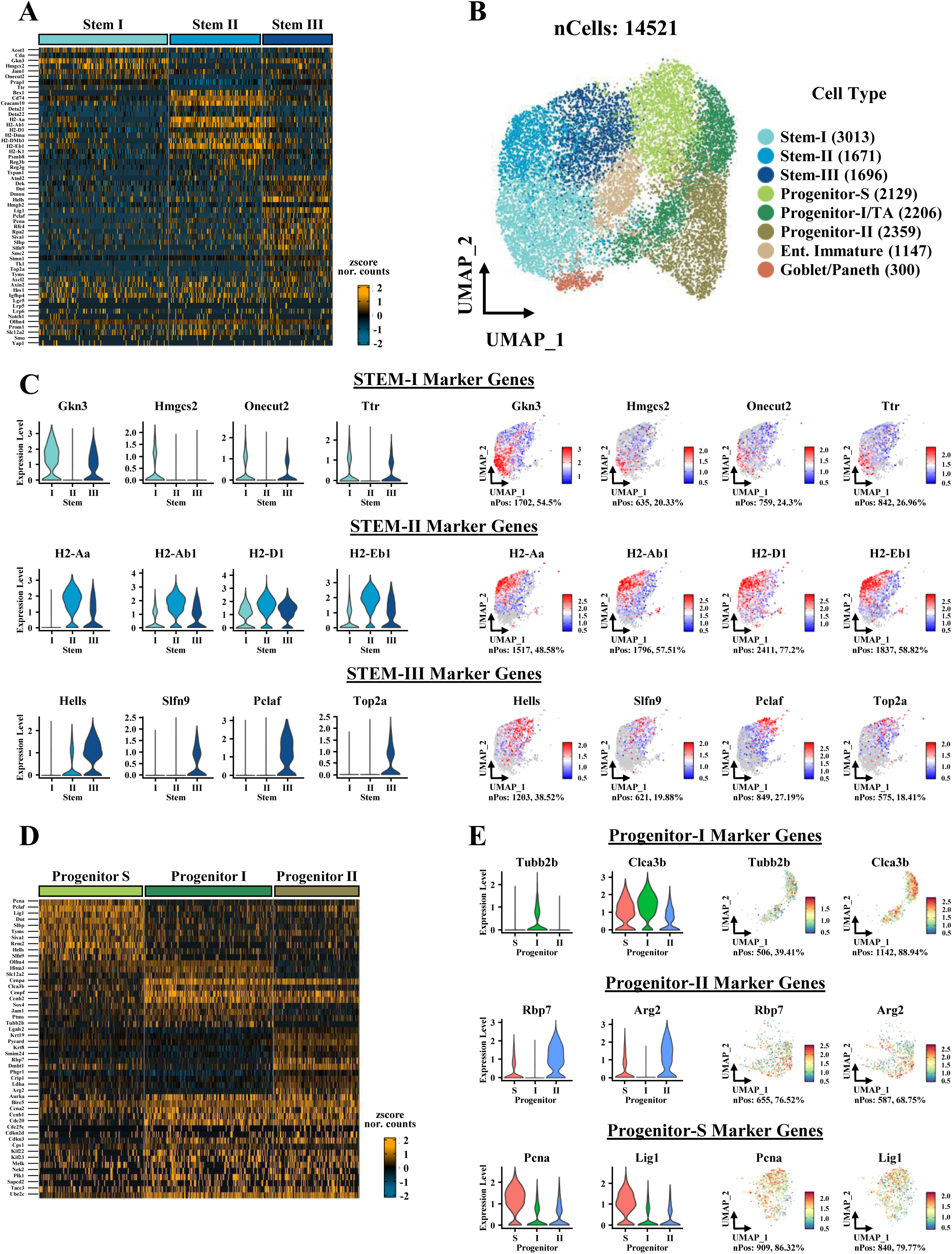
Gene signatures characterizing Stem cell and Progenitor cell subsets (A) Heatmap of relative mRNA levels (zscore normalized counts) of genes in Stem-I, Stem-II and Stem-III cell clusters. (B) UMAP projection of Stem cell and Progenitor cell types after sub-clustering of the indicated cell types. The color codes of the resulting re-clustered cell types are shown at the right. (C) Violin plots and Feature plots of selected marker genes characterizing Stem cell subpopulations. (D) Heatmap of relative mRNA levels (zscore normalized counts) of genes in Progenitor-I and Progenitor-II cell clusters. The gene expression profile of Progenitor S cells indicate that this cluster represents Progenitor cells in the S phase of the cell cycle. (E) Violin plots and Feature plots of selected marker genes characterizing Progenitor cell subpopulations.

**Figure S4.**
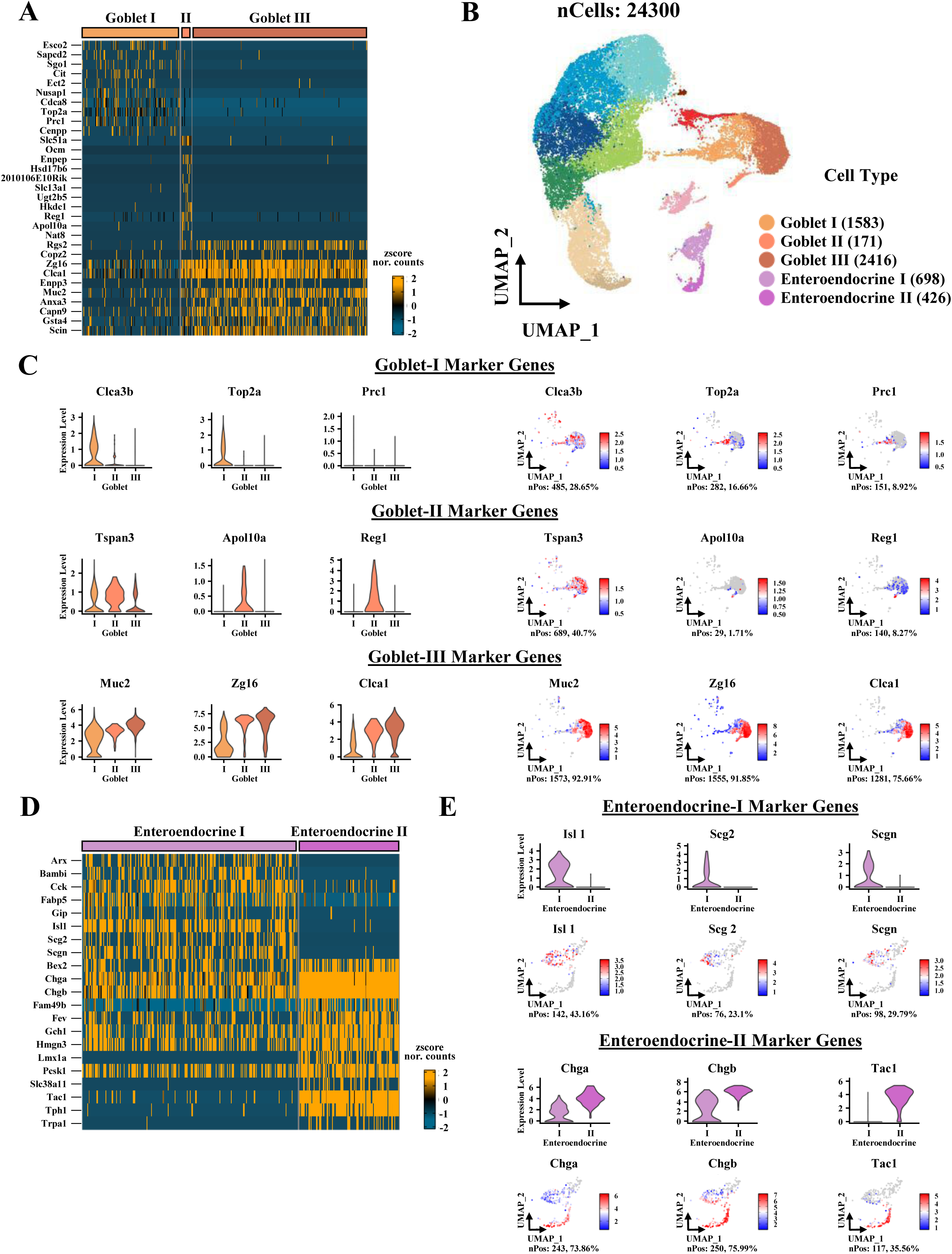
Gene signatures characterizing Goblet cell and Enteroendocrine cell subsets. (A) Heatmap of relative mRNA levels (zscore normalized counts) of genes in Goblet-I, Goblet-II and Goblet-III cell clusters. (B) UMAP projection of cell types from Figure S2A, used to characterize Goblet and Enteroendocrine cell subtypes. (C) Violin plots and Feature plots of selected marker genes characterizing Goblet cell subpopulations. (D) Heatmap of relative mRNA levels (zscore normalized counts) of genes in Enteroendocrine-I (EEC-I) and Enteroendocrine-II (EEC-II) cell clusters. (E) Violin plots and Feature plots of selected marker genes characterizing Enteroendocrine cell subpopulations.

**Figure S4.**
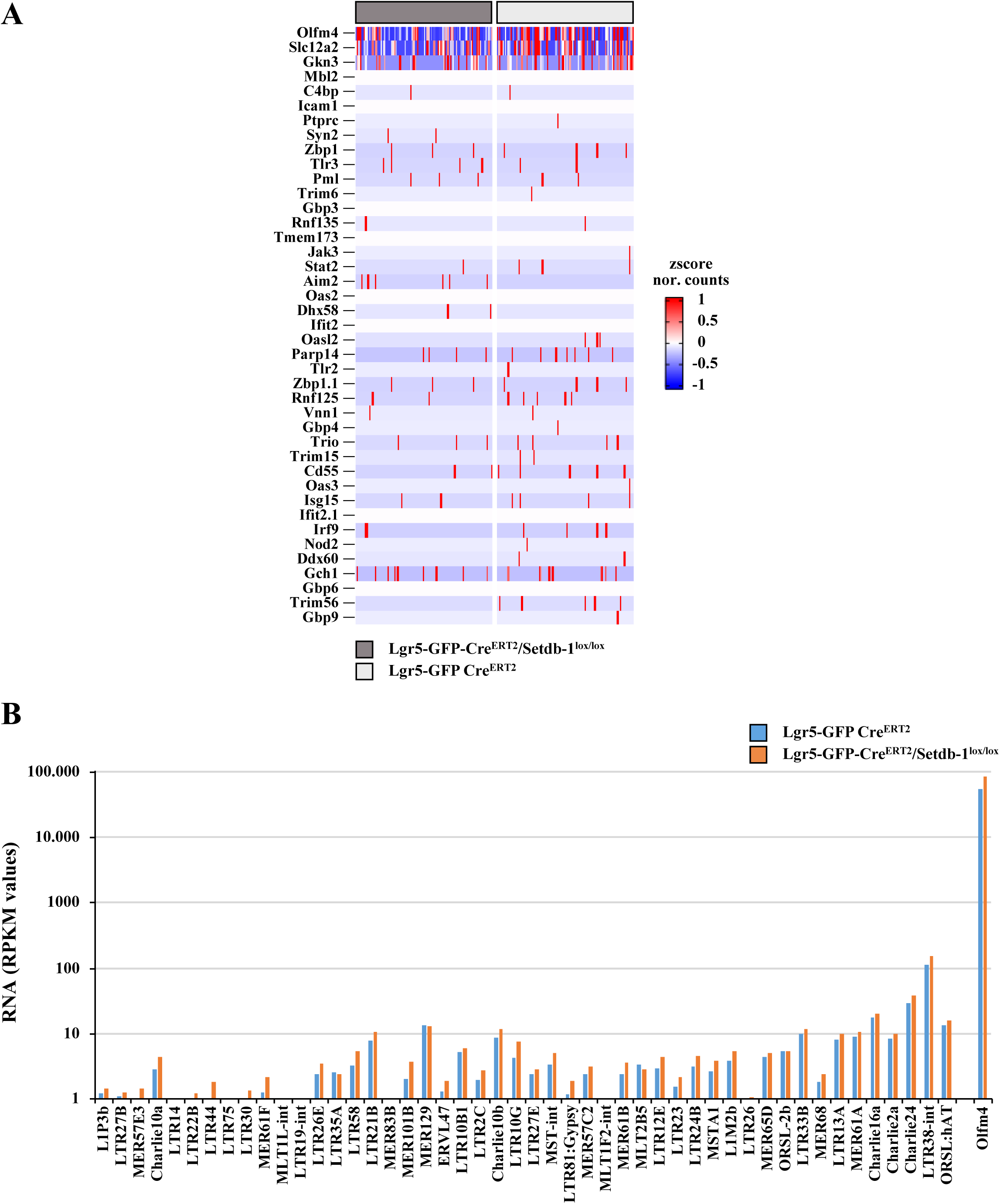
Lack of activation of innate immunity genes and endogenous retrovirus activation in Lgr5-GFP-Cre^ERT2^/ Setdb1^lox/lox^ cells 5 days after tamoxifen induction. (A) Heatmap showing RNA levels of innate immunity genes in individual cells from tamoxifen-treated Lgr5-GFP-Cre^ERT2^ (WT, light gray) and Lgr5-GFP-Cre^ERT2^/ Setdb1^lox/lox^ (Setdb1-KO, dark gray) mice. Heatmap of the highly expressed Olfm4 is included in the top panel. (B) Bar-graph showing relative mRNA levels of the indicated endogenous retroviral transcripts in FACS-sorted GFP^+^ cells from tamoxifen-treated Lgr5-GFP-Cre^ERT2^ mice and Lgr5-GFP-Cre^ERT2^/ Setdb1^lox/lox^ mice, as determined Tettranscript analysis of total RNA-seq data. Bars represent average RPKM values from two biological replicates. For comparison Olfm4 mRNA levels are included in the last bars. Note the 3 to 4 orders of magnitude difference between the retroviral RNAs and Olfm4 RNA levels and the small or no difference between the values obtained with wild type and Setdb1-deficient cells.

**Figure S6.**
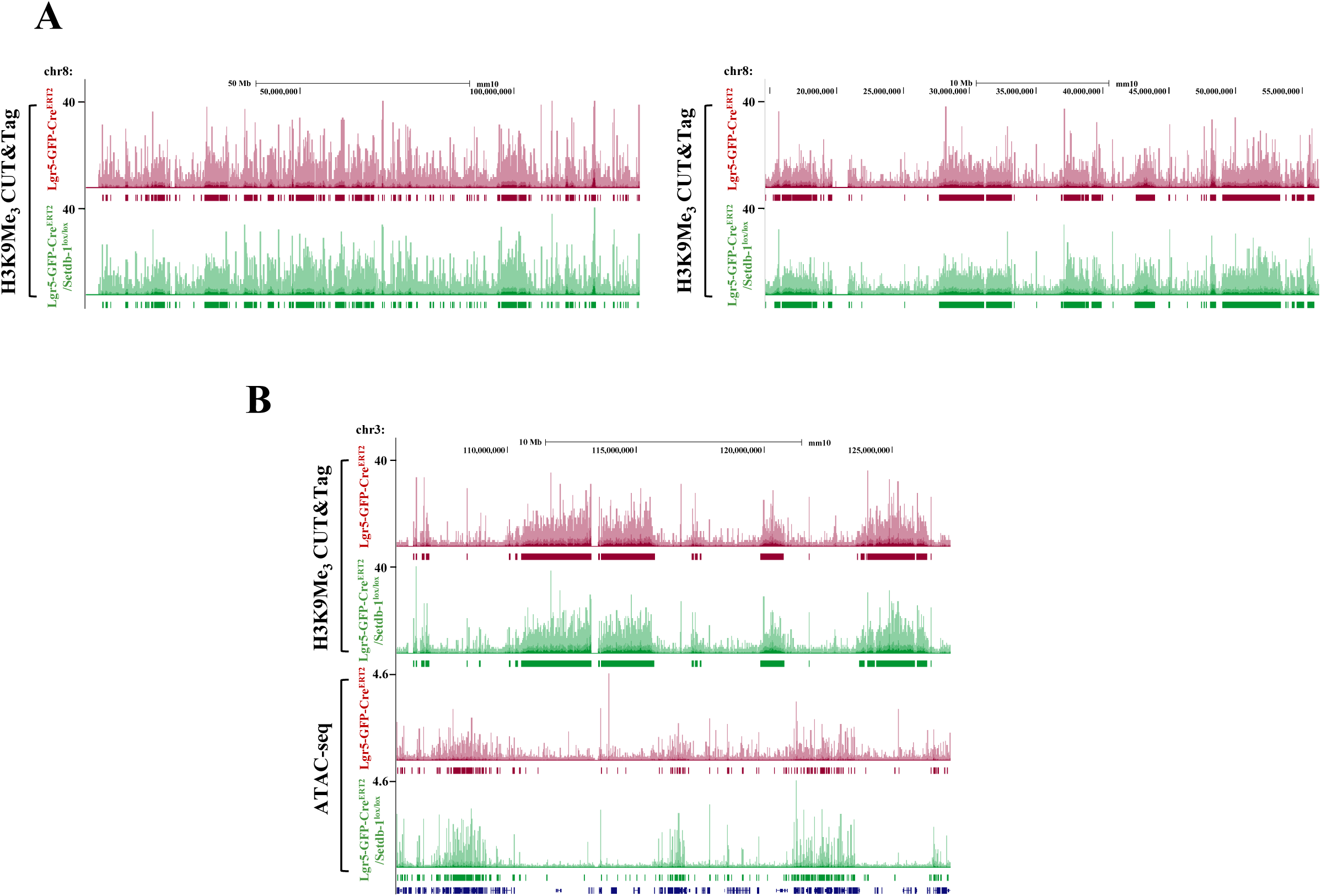
Limited changes in large H3K9Me_]_-modified heterochromatin domains in Setdb1-deficient cells. (A) Genome Browser tracks showing normalized H3K9Me_]_reads and SICER-called peaks (bars shown below the tracks) along the entire chromosome 8 (left panel) and a 40 Mb region of chromosome 8, from tamoxifen-treated Lgr5-GFP-Cre^ERT2^ mice (WT) and Lgr5-GFP-Cre^ERT2^/ Setdb1^lox/lox^ mice (Setdb1-KO). Note the highly similar overall distribution of the peak locations especially the broad peaks. (B) Combined Genome Browser tracks of H3K9Me_]_CUT&Tag and ATAC-seq reads spanning a 20Mb region of chromosome 3. Note the tandemly organized H3K9Me_]_-containing heterochromatin domains and the ATAC-seq signal containing euchromatin domains. Note the rare co-occurrence of ATAC-seq peaks and broad H3K9Me_]_CUT&Tag peaks in cells from Lgr5-GFP-Cre^ERT2^ mice, and their more frequent co-occurrence in Setdb1-KO cells.

**Figure S7.**
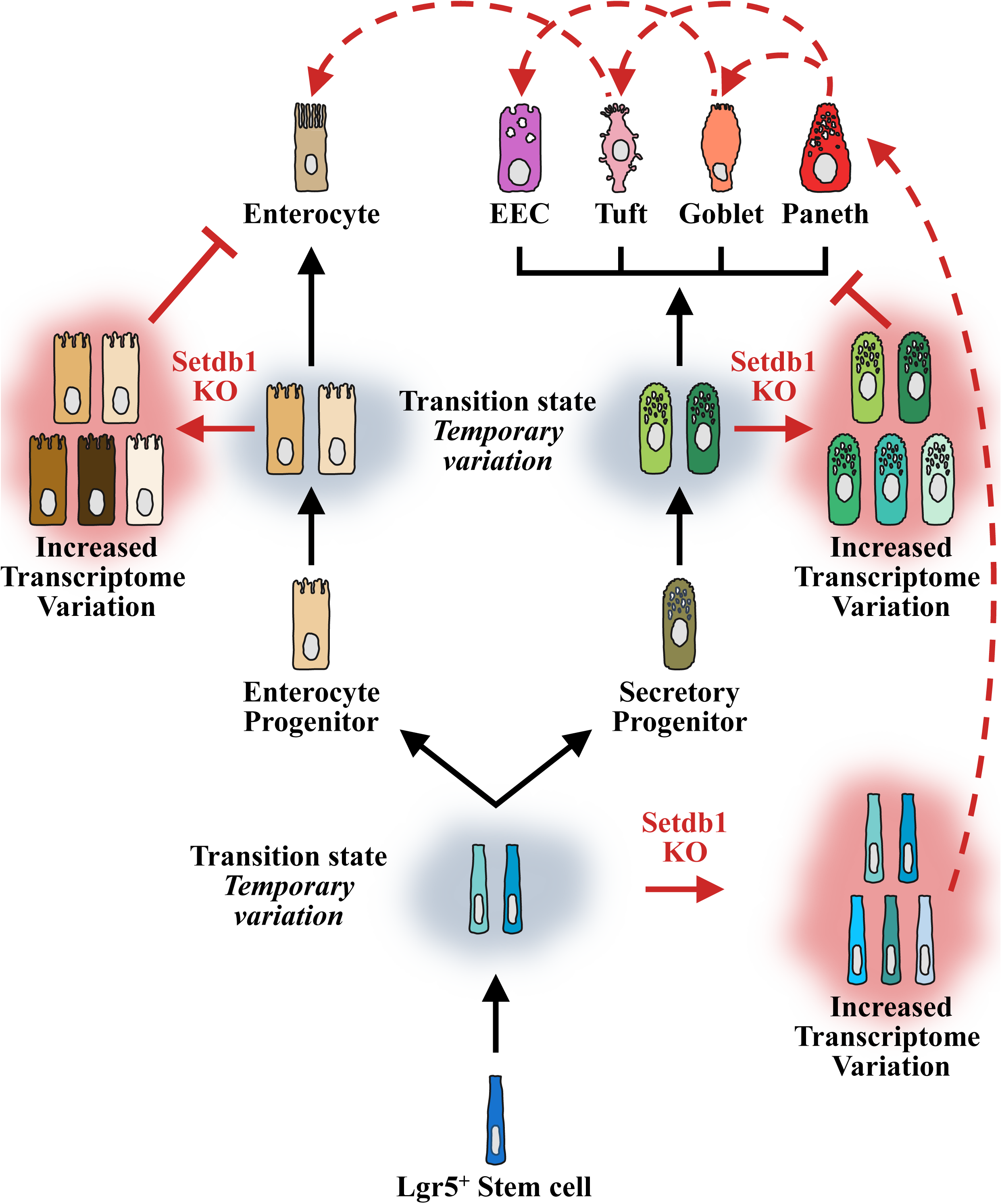
Model of chromatin defect-mediated increased cell-to-cell transcriptome variation during intestinal stem cell differentiation. The model proposes a multilevel control mechanism for the temporally distinct transcriptomes, at each consecutive differentiation state. Tight regulation promotes relatively uniform cellular transcriptomes, which is characteristic to Stem cells, Enterocyte Progenitor and Secretory Progenitors, as well as the differentiated epithelial cell types (Enterocytes, Paneth, Goblet, Tuft and Enteroendocrine cells). During the transition between the consecutive cellular differentiation states, partial transcriptome diversification occurs, leads to transition state-specific variable transcriptomes (cells in gray clouds). In Setdb1-KO mice, defects in chromatin structure-dependent regulation block cells in the transition states further increasing cell-to-cell transcriptional variations (cells in pink clouds), which cannot progress to fully differentiated cell types (flat-ended bar). Alternative pathways for the generation of secretory cell types directly from Stem cell population are triggered (dashed red arrow).

## Notes

### Competing Interest Statement

The authors have declared no competing interest.

### Summary of Updates

The revised version contains a more analysis of the single cell profiles. It has new data on transcription factor motifs and regulatory schemes associated with the loss of Setdb1. Figures 4 and 5 have been replaced by more accurate analysis and new data. New authors are added (E.D and G.G.) and authors order was partially changed.

